# Accuracy of *de novo* assembly of DNA sequences from double-digest libraries varies substantially among software

**DOI:** 10.1101/706531

**Authors:** Melanie E. F. LaCava, Ellen O. Aikens, Libby C. Megna, Gregg Randolph, Charley Hubbard, C. Alex Buerkle

## Abstract

Advances in DNA sequencing have made it feasible to gather genomic data for non-model organisms and large sets of individuals, often using methods for sequencing subsets of the genome. Several of these methods sequence DNA associated with endonuclease restriction sites (various RAD and GBS methods). For use in taxa without a reference genome, these methods rely on *de novo* assembly of fragments in the sequencing library. Many of the software options available for this application were originally developed for other assembly types and we do not know their accuracy for reduced representation libraries. To address this important knowledge gap, we simulated data from the *Arabidopsis thaliana* and *Homo sapiens* genomes and compared *de novo* assemblies by six software programs that are commonly used or promising for this purpose (ABySS, CD-HIT, Stacks, Stacks2, Velvet and VSEARCH). We simulated different mutation rates and types of mutations, and then applied the six assemblers to the simulated datasets, varying assembly parameters. We found substantial variation in software performance across simulations and parameter settings. ABySS failed to recover any true genome fragments, and Velvet and VSEARCH performed poorly for most simulations. Stacks and Stacks2 produced accurate assemblies of simulations containing SNPs, but the addition of insertion and deletion mutations decreased their performance. CD-HIT was the only assembler that consistently recovered a high proportion of true genome fragments. Here, we demonstrate the substantial difference in the accuracy of assemblies from different software programs and the importance of comparing assemblies that result from different parameter settings.

## Introduction

Advances in DNA sequencing have made the laboratory portion of large studies of genomic variation among many individuals feasible and economical, even for non-model taxa (Ekblom & Galindo, 2011; Narum *et al*., 2013; Andrews *et al*., 2016; Benestan *et al*., 2016). Current research in population genomics is often based on sequencing of reduced representation DNA libraries for many individuals, rather than the equivalent amount of whole genome re-sequencing for a smaller set of individuals (Buerkle & Gompert, 2013; Fumagalli, 2013). Ecological and evolutionary research in diverse systems often uses DNA sequencing to obtain genotypic information, regularly without a “complete” reference genome (draft genome assembly with long continguous sequences, for the focal or a related taxon). Instead, many studies rely on *de novo* assemblies of the subset of the genome contained in reduced representation libraries to use for reference-based mapping of reads. Despite the fact that the *de novo* assembly is prerequisite for further analysis and has strong potential to affect reference-based assemblies and genotyping, few studies have compared the efficacy of different software for sequence assembly from reduced representation libraries, including various protocols for genotyping-by-sequencing (GBS) and restriction associated DNA (RAD) sequencing (e.g., Baird *et al*., 2008; Elshire *et al*., 2011; Parchman *et al*., 2012; Peterson *et al*., 2012), which collectively we will refer to as GBS hereafter.

We performed a literature review to assess which *de novo* assembly programs are commonly used for processing GBS data. We report the software that was used in those studies and the extent to which papers evaluated and reported different assemblies based on different software settings and parameters. Some software was designed specifically for assembling RAD sequences (e.g., Stacks; Catchen *et al*., 2011; Rochette *et al*., 2019). Other programs were designed for assembly of whole genomes, transcriptomes, or protein sequences, including assembling contiguous sequences (contigs) that are longer than raw reads from the sequencing instrument (e.g., Velvet; Zerbino & Birney, 2008). These assemblers are sometimes applied to reduced representation sequences despite being designed for a different type of data, and the appropriateness of this application has not been directly evaluated.

There are a variety of parameter options for a given assembly method. For example, the assembly module in the software Stacks allows users to vary over a dozen parameters, whereas the software CD-HIT requires users to define only a few parameters (Catchen *et al*., 2011; Li & Godzik, 2006). Although assembly parameters such as percent match (i.e., threshold required for reads to be clustered into a single contig) can significantly alter resulting assemblies (Harvey *et al*., 2015; Ilut *et al*., 2014), few studies report any attempts to optimize these parameters. Resources to compare assemblies with different parameter settings and optimize assembly performance are increasingly available, but so far are underutilized (Paris *et al*., 2017; Puritz *et al*., 2014; Harvey *et al*., 2015; Ilut *et al*., 2014).

In this study, we evaluated the performance of a sample of leading *de novo* assembly programs. We used simulations of sequence reads for a population of individuals to compare *de novo* assembly software programs in terms of the accuracy of the resulting assemblies, given known locations for sequence reads within each genome. As representative, well-assembled genomes, we used the human (*H. sapiens* hereafter; GRCh38 retrieved from genome.ucsc.edu in Jan. 2014, Lander *et al*., 2001) and *Arabidopsis thaliana* (*A. thaliana* hereafter; TAIR 10 assembly, Lamesch *et al*., 2012) genomes and their expected fragmentation through double digestion with two restriction enzymes (EcoRI and MseI). We simulated different rates of mutation and different mutation types (single nucleotide polymorphisms and insertions/deletions) to evaluate assembler performance with different genome characteristics. Additionally, we investigated the sensitivity of the assemblers to two parameters that are used in their algorithms: percent match and k-mer length. We compared the six assemblers by quantifying different types of errors in their assemblies of our simulated data. We compared the completeness of the assemblies (fraction of all true genome fragments represented) and their degree of over-assembly (i.e., collapsing multicopy, paralogous loci into a single contig) and under-assembly (i.e., separating allelic variants at a single locus into different contigs).

## Methods

### Literature review

We performed a literature search using the Web of Science database to quantify the frequency of use of different assemblers in current GBS studies. Our search terms were “double digest” or “genotyping by sequencing” or “restriction site-associated”. We limited the search to papers from 1 January 2012 to 18 September 2017, a period that is relevant to GBS methods and generated an adequate sample size. We retained papers that presented new GBS data, performed a library preparation method that included digestion with two enzymes, and performed *de novo* assembly. We excluded single enzyme GBS studies because the assembly of single-digest fragments is a more complex problem, in part because their sequence reads are of either DNA strand surrounding the restriction site. We reviewed the papers in reverse chronological order of publication date and retained the first 100 papers that met these criteria to evaluate a reasonable subset of the relevant literature. For these 100 papers, we documented the *de novo* assembly software that was used and whether the authors varied assembler parameters, including percent match, k-mer length, or any other parameter.

### Assembler selection

We selected six software programs for assessment of their performance: ABySS, CD-HIT, Stacks, Stacks2, Velvet, and VSEARCH. We chose a sample of assemblers that were presented in peer-reviewed publications, employed a variety of assembly strategies, and were freely available. Additionally, assemblers were only included in the assessment if they had adequate and up-to-date user resources available online (e.g., manual, tutorial, user help forums). Lastly, we included assemblers that were commonly used in the published literature, and therefore likely of interest to researchers currently performing *de novo* assembly. The six assembly programs we selected represent variations on two clustering algorithms (graph-based and greedy clustering algorithms), and together these programs were used in 55% of the papers in our literature review. Although our comparison is not a complete list of assemblers meeting the desired criteria, our aim was to investigate variation in performance among a sample of currently available software.

We included four assemblers that use **graph-based algorithms** in our comparison: Stacks (version 1.46), Stacks2 (version 2.1), ABySS (version 1.3.4) and Velvet (version 1.1) (Catchen *et al*., 2011; Rochette *et al*., 2019; Simpson *et al*., 2009; Zerbino & Birney, 2008). We evaluated both Stacks and Stacks2 due to significant changes in the software related to how insertion and deletion (indel) variation is treated (changes to Stacks2 since the version 2.1 we used have not included changes to *de novo* assembly). These four assemblers apply graph theory, whereby nodes represent unique reads and edges connect nodes that have sequence segments in common. Graphs are constructed using maximum likelihood to cluster reads into contigs (Catchen *et al*., 2011). All four assemblers rely on input parameters to vary assembly constraints, but little guidance exists for their use with GBS data, except for efforts to aid users in selecting parameters for Stacks and Stacks2 (Paris *et al*., 2017). Because each assembler includes some unique parameters, we set parameters that we were not explicitly testing to comparable values when possible. For example, in Stacks and Stacks2 we set the minimum depth of coverage required to create a contig at 1 to mimic other assemblers having no rule for minimum requirement and performed an un-gapped alignment. We also allowed assemblers to optimize parameters when the option was available. Stacks, Stacks2, and a script for Velvet (VelvetOptimizer) were used to optimize k-mer length (Gladman & Seeman, 2012). VelvetOptimiser is substantially more memory intensive than simply running Velvet so we were unable to use it for the *H. sapiens* simulations. We chose to include assemblers designed specifically for reduced representation datasets (i.e., Stacks, Stacks2), as well as assemblers designed for other applications that are sometimes utilized for reduced representation datasets (i.e., for whole genome assembly using short reads: ABySS, Velvet).

We included two assemblers, CD-HIT (version 4.6.6) and VSEARCH (version 2.4.0), that use **greedy clustering algorithms** for assembly (Li & Godzik, 2006; Rognes *et al*., 2016). Greedy clustering algorithms group reads into clusters incrementally by optimizing similarity within contigs and dissimilarity between contigs. Although VSEARCH was intended for *de novo* assembly of metagenomic sequence data, it is also used for alignment and clustering in PyRAD, a program commonly used in GBS studies (Eaton, 2014). CD-HIT was developed for assembling protein sequences, but was later extended for nucleotide sequences, and it is used in the analysis pipeline dDocent (Puritz *et al*., 2014). We used dDocent’s data reduction step that retains only one copy of each unique sequence for assembly to reduce computational time (this script can be found in the Dryad repository: (https://doi.org/10.5061/dryad. 8tr03f8, LaCava *et al*.)).

### Parameter settings

We varied two parameters, percent match and k-mer length, across all assemblers and simulations to evaluate their influence on assembly. We selected 90%, 94%, and 98% for the minimum percent match to investigate how these interacted with allelic and paralogous variation in the simulated reads. Some assemblers use percent match as a parameter (e.g., CD-HIT), while others use raw mismatch numbers (e.g., Stacks), so an apparently identical parameter setting could produce varied results depending on this distinction. For example, a 100bp read that includes a 6bp barcode results in 94bp reads at the assembly step. For our 98% match setting, the raw mismatch assemblers would allow 2 base pairs to mismatch, while the percent match assemblers would require 98% match, and 0.98×94bp = 92.12, which allows only one base pair to differ. This is consistent across all three percent match settings; for assemblers that use percent match, the mismatch allowed is one base pair less than for assemblers that use raw mismatch. Table 2 identifies the version of this parameter used in each assembler, and lists base pair equivalents of the percent match used.

**Table 1:** Parameter settings for *in silico* restriction enzyme digestions and simulated reads for GBS using ddRADseqTools (Mora-Márquez *et al*., 2017). The parameter mutprob sets the probability that a base pair in the locus will mutate. The maximum mutations allowed per locus is set by locusmaxmut. The probability that a mutation will be an insertion or deletion (indel) rather than a SNP is set by indelprob. The maximum length in base pairs of the indels is set by maxindelsize. We simulated reads for both the *A. thaliana* and *H. sapiens* reference genomes using each of these nine parameter combinations.

| Simulation Name | Description | `mutprob` | `locusmaxmut` | `indelprob` | `maxindelsize` |
| --- | --- | --- | --- | --- | --- |
| Original genome | No mutation in homozygous genome | 0 | 3 | 0 | 1 |
| A few SNPs | SNPs only, at a low probability | 0.1 | 3 | 0 | 1 |
| More SNPs | SNPs only, at high probability | 0.2 | 3 | 0 | 1 |
| Multiple SNPs per locus | High number of mutations per locus | 0.1 | 5 | 0 | 1 |
| Mostly SNPs, few indels | Most mutations are SNPs, 10% indels | 0.1 | 3 | 0.1 | 1 |
| 50:50 SNPs:indels | Half of mutations are SNPs, half indels | 0.1 | 3 | 0.5 | 1 |
| Only 1 bp indels | Mutations are mostly 1 bp indels | 0.1 | 3 | 0.99 | 1 |
| 1–3 bp indels | Mutations are mostly 1–3 bp indels | 0.1 | 3 | 0.99 | 3 |
| 1–5 bp indels | Mutations are mostly 1–5 bp indels | 0.1 | 3 | 0.99 | 5 |

**Table 2:** Percent match and k-mer length values tested for each assembler. We tested a range of parameter values possible for each assembler. We also constructed assemblies using the assembler-optimized parameter values, or if the assembler did not have an optimization routine, we used the defaults. Optimized or default parameter values are in bold. For the match parameter, assemblers either use raw mismatch or percent match, and with 94bp reads this means the same parameter values result in different raw mismatch values, indicated here. *Note that for Velvet, only the *A. thaliana* simulations had optimized k-mer length because the attempted optimization of *H. sapiens* simulations exceeded available computational resources.

| Assembler | K-mer length | Match parameter | Raw mismatch allowed |
| --- | --- | --- | --- |
| ABYSS | 15, 31, **optimized** | Raw mismatch | 10, 6, 2 |
| CD-HIT | NA | Percent mismatch | 9, 5, 1 |
| Stacks | 15, 31, **optimized** | Raw mismatch | 10, 6, 2 |
| Stacks2 | 15, 31, **optimized** | Raw mismatch | 10, 6, 2 |
| Velvet | 15, 31, **optimized\*** | Percent mismatch | 9, 5, 1 |
| VSEARCH | **8 (default)**, 15 | Percent mismatch | 9, 5, 1 |

We compared k-mer lengths of 8–31bp, as well as an assembler-optimized k-mer length, although some assemblers were unable to run with every k-mer length (Table 2). K-mer length represents the sequence length that the assembler algorithm uses to compare reads; that is, the algorithms do not consider the entire, intact sequence at once.

### Simulations

As representative genomes, we selected the *A. thaliana* (1.44 × 10^8^ base pairs, including gaps and unknown bases, unambiguously mapped to chromosomes, mtDNA, and cpDNA in TAIR10) and *H. sapiens* (3.85 × 10^9^ base pairs, including gaps and unknown bases, unambiguously mapped to chromosomes and mtDNA in GRCh38) genomes to evaluate to what extent genome complexity influences assembler performance. These genomes differ in total genome size and amount and structure of the repetitive content in their genomes, potentially presenting different challenges for *de novo* assembly.

We used ddRADseqTools version 0.42 (https://github.com/GGFHF/ddRADseqTools, Mora-Márquez *et al*., 2017) to create *in silico* ddRAD digests of the *A. thaliana* and *H. sapiens* genomes. From the ddRADseqTools package, we used the script *rsitesearch.py* to obtain fragments from restriction sites for the commonly used EcoRI and MseI restriction enzymes within each of the reference genomes. We then used the script *simddradseq.py* to set parameters and simulate 100 bp single-end reads, including barcodes, from the genome fragments. Because we were not interested in investigating additional complexity introduced by PCR error, we did not simulate PCR duplicates and did not use the PCR duplicate removal step in *pcrdupremoval.py*. We used *indsdemultiplexing.py* to demultiplex the simulated reads and *readstrim.py* to trim the barcodes from the reads, resulting in reads of 94 bp that all began with the EcoRI restriction site. Our wrapper functions for each simulation, modified from *run ddradseq chain.sh* in ddRADseqTools, are included in the Dryad repository: (https://doi.org/10.5061/dryad.8tr03f8, LaCava *et al*.).

We used the *in silico* ddRAD digests of *A. thaliana* and *H. sapiens* genomes to simulate sets of reads for each genome. We simulated 100 individuals for all simulations of both genomes. We set minfragsize = 350 and maxfragsize = 400 to simulate size selection of fragments between 350 and 400 bp (in practice, a larger range of fragments would mean more paralogous loci would be in the sequenced library). We set minreadvar = 1 and maxreadvar = 1 for all simulations so that read depth was approximately constant across all fragments. We set locinum to the number of fragments obtained by *rsitesearch.py* for each reference genome, and then set readsnum to a sufficiently high number that all fragments were sampled in the output reads. For *A. thaliana* we set locinum = 1,849 and readsnum = 3,698,000; for *H. sapiens* we set locinum = 45,190 and readsnum = 9,0380,000.

We generated nine sets of simulated reads per genome with varying rates and types of mutations (Table 1). We varied mutation probability, maximum number of mutations per locus, mutation type (i.e., probability of nucleotide variation [SNP], or nucleotide insertion or deletion [indel]), and maximum mutation length (for indels). Mutations based on these parameters are randomly introduced within simulated reads using the Jukes-Cantor model of sequence evolution (assumes equal nucleotide frequencies and equal mutation rates across base pairs), and the location of mutations is conserved across loci and individuals within each simulated dataset. The first simulation had mutprob set to zero, so this simulation recovered the original genome. The other eight simulations for each reference genome varied in the amount and types of mutations. We refer to the 350–400 bp reference genome sequences as ‘fragments’, sequences produced by the simulator from the fragments as ‘reads’, and sequences determined by the assemblers to be unique parts of the reference set as ‘contigs’.

### Quantifying accuracy of assemblies

We constructed GBS assemblies using each assembler for the nine simulated data sets for *A. thaliana* and *H. sapiens*. We used a custom Perl script to compare the assembled contigs with the known fragments from the in-silico digestion of genomes to determine assembly accuracy using two metrics (this script can be found in the Dryad repository: (https://doi.org/10.5061/dryad.8tr03f8, LaCava *et al*.)). To evaluate how completely each assembler recovered the original genome fragments (loci), we counted the number of true genome fragments that were represented in the assembly (*completeness* criterion). We used the simulation’s record of what genome fragment a simulated read had been drawn from (recorded in the information line of reads in the simulated fasta data), regardless of whether the read corresponded to the ancestral or a mutated sequence. For the completeness criterion, we counted both assembled contigs that perfectly matched simulated sequences, as well as contigs that were at least 94 base pairs in length and contained the full length of a simulated sequence, but potentially contained additional bases to accommodate indel variation. Assemblies would be incomplete if some fraction of true fragments and their corresponding ancestral or mutated sequences were not represented in the contigs, because true fragments had been incorrectly subdivided and shortened.

Furthermore, whereas an assembly could be a complete representation of all fragments in the genome, those fragments could be under- or over-assembled relative to the true number of unique genomic regions. Thus, we also compared the number of true fragments in the genome to the number of assembled contigs (*over-under assembly* criterion). A correct assembly would produce exactly as many contigs as there are unique fragments, regardless of mutations. An under-assembled genome would contain more contigs than there are fragments (i.e., fragments incorrectly split into more contigs than is accurate; a ratio greater than one), whereas an over-assembled genome would contain fewer contigs than there are fragments (i.e., fragments collapsed into fewer contigs than is accurate; a ratio less than one). Over- and under-assembly can significantly affect downstream analyses, and while some assembly errors can be identified and accounted for in subsequent filtering steps, the reliance on filtering to remove assembly errors is not ideal. Therefore, software that avoids both over- and under-assembly is preferable.

## Results

### Literature review

We reviewed a total of 665 papers to find 100 papers that met the desired criteria. The 100 selected papers spanned February 2015 to October 2017. Of the 100 papers, 39 used Stacks (the period we reviewed preceded the release of Stacks2), 19 used UNEAK, 11 used VSEARCH, and 14 used one of the following assemblers: DNASTAR SeqMan, dDocent (i.e., CD-HIT), or AftrRAD. The remaining 17 papers each used a unique assembler (see Table S1). Of 100 papers, 13 reported that they varied percent match, while 12 reported that they varied other assembler parameters. None of the reviewed papers reported varying k-mer length.

### Simulations

The *A. thaliana* genome contained 1849 GBS fragments in the 350–400 bp size class, but when we simulated 100 bp reads from these fragments and removed the 6 bp barcode from the beginning of the read, this reduced to 1813 unique DNA sequences. The *H. sapiens* genome contained 45190 fragments in the 350–400 bp size class, corresponding to 43160 unique sequences when trimmed to 94 bp. Since *de novo* assemblers cannot distinguish identical sequences from different parts of the genome, we used the unique 94 bp fragments to represent the expected number of contigs in our analyses of assembler performance.

### Recovery of genomes without simulated mutation

Across all k-mer length and percent match settings, CD-HIT, Stacks, Stacks2, and VSEARCH recovered at least 96% of the fragments from the unmutated *A. thaliana* genome (Figure 1, Table S2). Velvet assemblies recovered 83–95.0% of the fragments from the unmutated *A. thaliana* genome, depending on the k-mer length setting (Figure 1). In the larger and more complex *H. sapiens* genome, CD-HIT, Stacks, Stacks2 and VSEARCH recovered at least 87% of the fragments from the unmutated genome across all percent match and k-mer length settings (high completeness; Figure 1). Of all assemblers, CD-HIT with a percent match setting of 98% recovered the highest proportion of fragments (98.3%) from the unmutated *H. sapiens* genome (Table S3). Velvet recovered 17% (k-mer length=15) to 76% (k-mer length=31) of the fragments from the unmutated *H. sapiens* genome, regardless of percent match setting (Figure 1). ABySS failed to recover any full-length contigs that corresponded to fragments from the unmutated *A. thaliana* and *H. sapiens* genomes (Tables S2 & S3). Instead, ABySS retained contigs that corresponded to fragmented sequence reads, and reported a large number of ”contigs” that were shorter than, and lost information relative to, the simple set of unique sequence reads. Thus, we excluded ABySS from further analysis.

**Figure 1:**
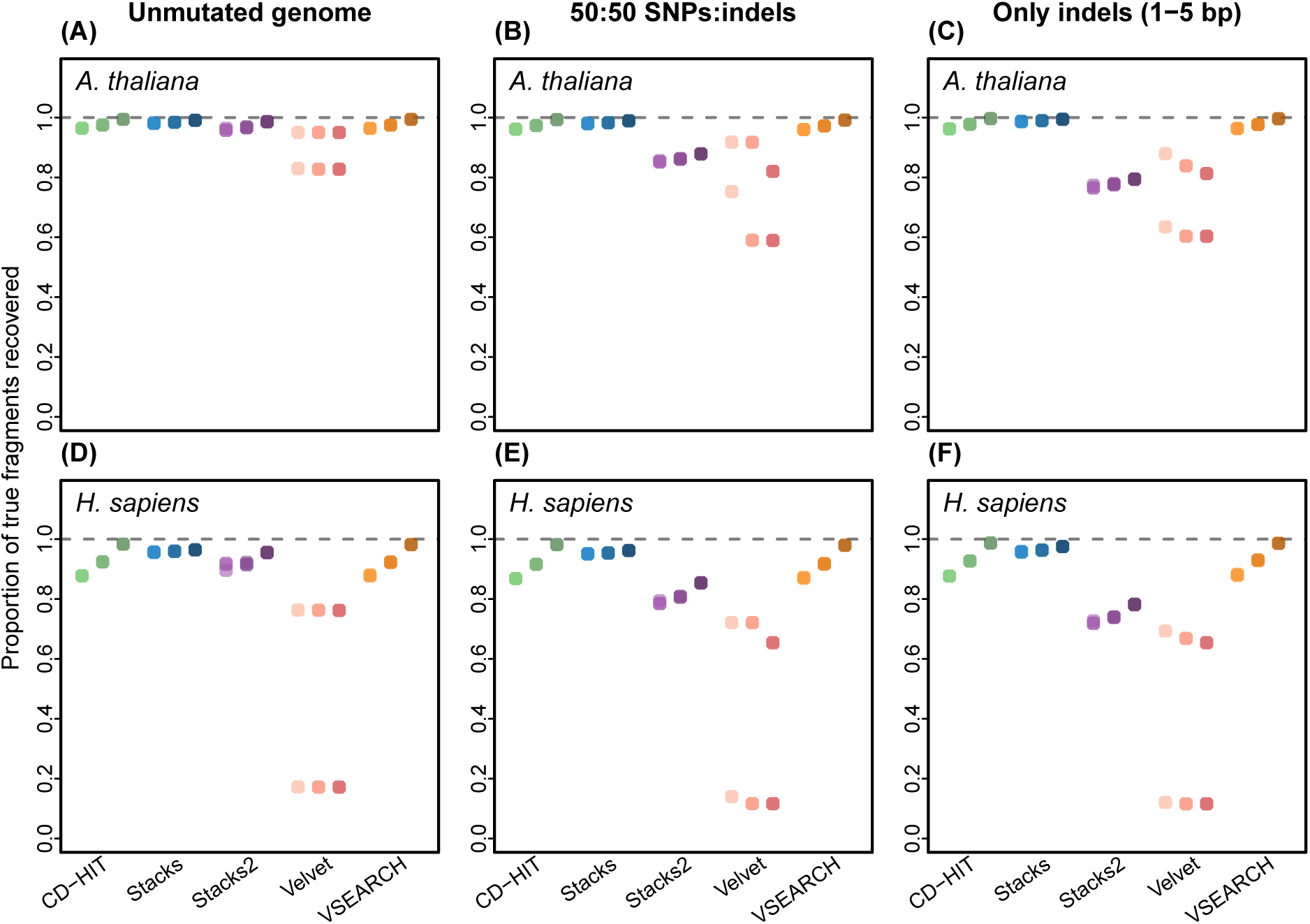
The completeness of assemblies in simulations of unmutated genomes (A, D), in simulations of an equal number of SNPs and indels (B, E), and simulations of 1–5 base pair indels (C, F). Simulations were derived from the *A. thaliana* (A-C) and *H. sapiens* (D-F) genomes. Completeness was calculated as the proportion of contigs that matched original genome fragments. A value less than 1 indicates that some of the assembled contigs were not found in the genome fragments. Values are reported for five assemblers: CD-HIT (green), Stacks (blue), Stacks2 (purple), Velvet (pink) and VSEARCH (orange). The hue of each color corresponds to the percent match parameter setting used in the assembly, with light hues corresponding to 90% match, medium hues corresponding to 94% match, and dark hues corresponding to 98% match. Assemblers have multiple dots in the same hue when k-mer length affected assembly outcome (see Tables S2 & S3 for details).

**Table 3:** A summary of assembler performance. Assemblers are compared based on recovery of the original genome, assembly outcomes when presented with simulated data with differing degrees of mutations arising from SNPs and indels, and the impact of varying the k-mer and percent match parameter settings.

| Assembler | Genome recovery | Assembly of SNPs | Assembly of indels | Sensitivity to k-mer | Sensitivity to % match |
| --- | --- | --- | --- | --- | --- |
| ABYSS | Worst | NA | NA | NA | NA |
| CD-HIT | Best | Best | Best | NA | Sensitive |
| Stacks | Best | Best | Poor | Relatively insensitive | Relatively insensitive |
| Stacks2 | Best | Best | Poor | Relatively insensitive | Relatively insensitive |
| Velvet | Poor | Mixed | Mixed | Very sensitive | Sensitive |
| VSEARCH | Mixed | Worst | Worst | Very sensitive | Sensitive |

None of the assemblers recovered the exact number of contigs expected. CD-HIT, Stacks, Stacks2 and Velvet over-assembled contigs to varying degrees (Figure 2). Assemblies from CD-HIT, Stacks and Stacks2 recovered 88-98% of true fragments (contig/fragment ratios), whereas Velvet assemblies contained 77% (with k-mer length of 31) to 93% of the true number of fragments (k-mer length of 15), regardless of percent match setting. VSEARCH was the only software that under-assembled contigs when assembling the original, unmutated genomes (Figures S1 & S2). For VSEARCH, a k-mer length of 8 resulted in approximately the correct number of contigs, but a k-mer setting of 15 resulted in under-assembly of the *A. thaliana* genome, producing two times more contigs than the true number of unique fragments (Table S2). Under-assembling was more severe for VSEARCH with the more complex *H. sapiens* genome, resulting in over three times the number of contigs compared to the expected number (Table S3).

**Figure 2:**
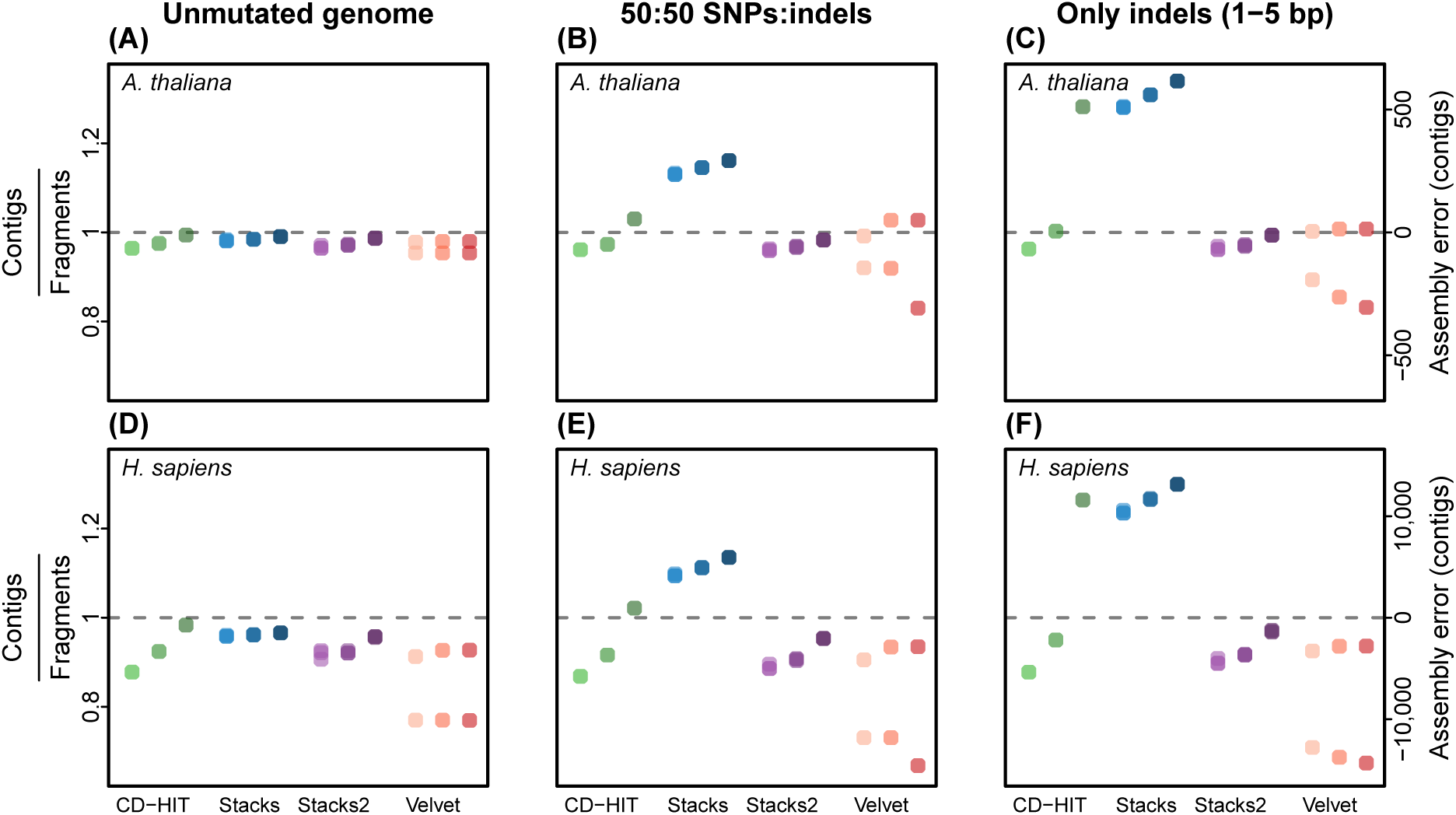
Measures of over- and under-assembly in simulations of unmutated genomes (A, D), in simulations of an equal number of SNPs and indels (B, E), and simulations of 1–5 base pair indels (C, F). Simulations were derived from the *A. thaliana* (A–C) and the *H. sapiens* (D–F) genomes. Over- and under-assembly are presented by the ratio of assembled contigs to true genome fragments (left vertical axis) and by absolute numbers (right vertical axis). A contigs:fragments ratio greater than one represents under-assembly and a ratio less than one represents over-assembly. Assembly results are shown for CD-HIT (green), Stacks (blue), Stacks2 (purple) and Velvet (pink) with variable percent match (light hues = 90%, medium hues = 94%, dark hues = 98%). Assemblers have multiple dots in the same hue when k-mer length affected assembly outcome (see Tables S2 & S3 for details). VSEARCH was omitted because its under-assembly was so much greater than other software (VSEARCH is included in Figures S1 & S2).

### Sensitivity to SNPs

Across all simulations containing only SNPs, CD-HIT, Stacks, and Stacks2 recovered a high proportion of true genome fragments (Figures S3 & S4) and produced approximately the expected number of contigs (Figures S1 & S2). CD-HIT under-assembled SNP simulations when we used a percent match parameter setting of 98%, and the magnitude of under-assembly was greatest for simulations that allowed up to 3–5 SNPs per locus (Figures S1 & S2), where variation in the simulated reads was higher than variation permitted by the percent match setting. Across all SNP simulations, Velvet assemblies with a k-mer length of 15 resulted in over-assembly (Figures S1 & S2), whereas a k-mer length of 31 resulted in more accurate assemblies, similar to those produced by CD-HIT, Stacks, and Stacks2 (Figures S1 & S2). VSEARCH assemblies of simulated *A. thaliana* data varied from slight over-assembly to considerable under-assembly depending on k-mer length and the probability of SNPs in the simulated dataset (Figure S1). VSEARCH assemblies of simulated *H. sapiens* data containing SNPs consistently resulted in under-assembly (Figure S2).

### Sensitivity to indels

Across all three simulations that introduced only indels as mutations (Table 1), CD-HIT consistently recovered at least 96% of *A. thaliana* fragments and at least 87% of *H. sapiens* fragments (Figures S3 & S4).

CD-HIT only under-assembled with a percent match of 98% when indels were up to 3– 5 base pairs in length, where variation in the simulated reads was higher than variation permitted by the percent match setting (Tables 1, S2 & S3). Stacks consistently recovered at least 95% of true fragments from both genomes, but Stacks also consistently under-assembled, with contig/fragment ratios over 1.2 (Figures S1, S2, S3, & S4). This means that although all the contigs originated from true genome fragments, more contigs were produced than expected. In contrast, Stacks2 recovered only 71–82% of true genome fragments, but produced contig/fragment ratios of 0.90–0.97, meaning that Stacks2 produced closer to the correct number of contigs, but fewer of these contigs were found in the simulated genome fragments (Figures S3 & S4).

Similar to SNP simulations, Velvet assemblies for indel simulations varied in accuracy across k-mer settings. A k-mer setting of 15 produced approximately as many contigs as expected, but as few as 12% of those contigs were found in the original fragments. In contrast, a k-mer setting of 31 produced a contig/fragment ratio as low as 0.72, but a higher percentage of contigs matched true genome fragments (Figures S1, S2, S3, & S4). As with SNPs, VSEARCH performance varied between the *A. thaliana* and *H. sapiens* genomes. For *A. thaliana*, VSEARCH varied from slight over-assembly to considerable under-assembly depending on k-mer length and the length of indels simulated (Figure S1). Similar to SNP simulations, all indel simulations for *H. sapiens* resulted in under-assembly when using VSEARCH (Figure S2).

### Sensitivity to the combination of SNPs and indels

For assemblies of sequences that contained a combination of SNPs and indels (Table 1), CD-HIT, Velvet and VSEARCH performed similarly to simulations where SNPs or indels were introduced independently (Figures S1 & S2). The performance of Stacks with a combination of SNPs and indels, however, produced results intermediate to SNPs or indels independently. In the simulation with mostly SNPs and a few indels (Table 1), Stacks produced approximately the expected number of contigs (Figures S1 & S2). In the simulation with 50:50 SNPs and indels, Stacks consistently under-assembled for both genomes and across percent match settings (Figure 2). The degree of under-assembly by Stacks increased as more indels were introduced to simulations (Figure 2).

Stacks2 assemblies were also intermediate for simulations with a combination of SNPs and indels compared to simulations with each mutation type independently. As the proportion of indels increased, Stacks2 assemblies for both *A. thaliana* and *H. sapiens* were less complete, but the ratio of contigs to fragments remained close to 1 (Figures 1 & 2).

### Sensitivity to k-mer setting

K-mer length had almost no effect on the proportion of true genome fragments recovered or the number of contigs produced by Stacks or Stacks2 (Tables S2 & S3). However, k-mer length affected assembly outcome for both Velvet and VSEARCH across all simulations that included mutations (Figures S1, S2, S3, & S4). For Velvet, k-mer length affected both the proportion of the true fragments recovered and the rate of over-assembly. Across all simulations for both *A. thaliana* and *H. sapiens* genomes, a k-mer length of 15 consistently reduced the completeness of assemblies when compared to assemblies of k-mer length of 31. In contrast, a k-mer length of 31 typically resulted in over-assembly from Velvet, which was more extreme in the complex *H. sapiens* genome. For VSEARCH, k-mer length had little effect on the rate of recovery of true fragments, but can lead to substantial under-assembly (Figures S1 & S2). CD-HIT does not permit users to vary k-mer length, so this parameter was not evaluated for that assembler.

### Sensitivity to percent match

Stacks and Stacks2 were the least sensitive to varying the percent match parameter setting (Tables S2 & S3). CD-HIT and VSEARCH assemblies were affected by the percent match parameter setting in an expected fashion; for example, increasing percent match to 98% resulted in increased under-assembly for simulations that produced reads that diverged from the original fragments by more than two base pairs (Figures S1 & S2). Velvet assemblies either varied little with the percent match parameter setting (in the case of k-mer lengths of 15) or varied in the opposite direction (in the case of k-mer lengths of 31); the 98% match often caused greater over-assembly than the 90% or 94% match settings (Tables S2 & S3).

## Discussion

With any short read sequencing technology (commonly 100–250bp), there is some ambiguity in the alignment or mapping of those reads because of sequence similarity due to paralogy or allelic variation (Harvey *et al*., 2015; Ilut *et al*., 2014). This applies to mapping reads to a high-quality reference genome (e.g., with bwa, Li *et al*., 2010), to the *de novo* assembly of reads as investigated here for GBS data, or to the related challenges for *de novo* assembly of transcriptomes. The ambiguity due to sequence paralogy is evident in the 1.9–4.5% of GBS loci from the *A. thaliana* and *H. sapiens* genomes that were not distinguishable using 94 bp (assuming 6 bp of the 100 bp reads were used as molecular barcodes to distinguish samples). In typical molecular ecology studies, the problem is compounded by allelic variation in the sample used to construct the *de novo* reference genome. Consequently, some methods and models for identifying variants and calculating genotype likelihoods use mapping quality of reads (e.g., FreeBayes; Garrison & Marth, 2012), or use filtering steps to remove sites with low mapping quality scores. Longer reads, or pairs of reads that together are both longer and potentially separated in the genome by some length, will have a higher chance of mapping uniquely, but also have a higher chance of containing nucleotide variants relative to the reference genome or the alleles of other individuals. Thus, the problem of correctly mapping sequences to a reference or *de novo* assembly is general and not restricted to GBS data. In this study we have focused on the assembly problem in the context of GBS, because of the method’s common usage (reviewed in Ekblom & Galindo, 2011; Narum *et al*., 2013; Andrews *et al*., 2016; Benestan *et al*., 2016) and its potentially attractive place in the trade-offs that exist among: 1) completeness of genome coverage (low for GBS, relative to whole genome sequencing [WGS]), 2) depth of sequencing at locus (can be optimized, potentially high relative to WGS; Buerkle & Gompert, 2013; Fumagalli, 2013), and 3) numbers of individuals (can be optimized, potentially high relative to WGS) for a finite amount of sequencing. Despite legitimate concerns about the adequacy of genome coverage by GBS-like methods for certain questions and in some systems (Lowry *et al*., 2016), for many applications in population genomics GBS-like methods are likely to remain attractive for some time (McKinney, 2016). Whereas several studies have examined the consequences of laboratory and bioinformatic methods for variant identification and other downstream analyses (Shafer *et al*., 2017; Flanagan & Jones, 2018; Warmuth & Ellegren, 2019), and others have suggested methods to optimize assembly parameters (Puritz *et al*., 2014; Paris *et al*., 2017), this investigation fills a gap in knowledge regarding the performance of *de novo* assembly software without the aid of additional steps.

Our literature review of 100 recently published papers indicates that Stacks has been the most commonly used *de novo* assembler for GBS data (39 of the reviewed studies), but also that a large variety of software programs are used. Our comparative simulation study showed that Stacks (and Stacks2) recovered true genomes well in the absence of allelic variation, but did less well than CD-HIT (used in only 4 of 100 reviewed papers) for both the *A. thaliana* and *H. sapiens* genomes when mutations were present (Table 3). In particular, insertion and deletion polymorphisms caused under-assembly of reads for Stacks (as previously demonstrated by Puritz *et al*., 2014) and a failure to recover a substantial fraction of true genome fragments for Stacks2 (presumably because polymorphisms led to fragmentation of contiguous sequences in the assemblies). CD-HIT was the only assembler that across simulations consistently recovered a high proportion of true genome fragments and its assemblies typically were close to the original genome fragments (with the expected exception in assemblies with a 98% minimum match percentage separating, in which allelic variants with greater than 2% divergence were placed into separate contigs). Two of the other assemblers we considered, Velvet (used in 1 of 100 reviewed papers) and VSEARCH (used in 11 of 100 reviewed papers), either performed relatively well at recovering all genome fragments, or at assembling reads into the correct number of genome fragments, but not both. Somewhat dependent on the k-mer setting and the simulation, Velvet assemblies failed to recover a substantial fraction of true fragments (sometimes with counter-intuitive sensitivity to assembler settings), yet over-assembled those fragments only modestly. Whereas VSEARCH assemblies recovered a high fraction of the true genome fragments across simulations, typically the assemblies were drastically under-assembled, particularly for the human genome. Finally, ABySS was poorly suited to *de novo* assembly of GBS reads (it was not used in any of the 100 reviewed studies), in that it resulted in contigs that were exclusively shorter than the original reads and its assemblies did not contain any true genome fragments.

We found that assembly method did not predict assembler performance in any consistent manner (Table 3). We included software that used either graph-based algorithms or greedy-clustering algorithms, and assemblers in each category varied in their performance. The highest performing assemblers, CD-HIT and Stacks2, used different algorithms, suggesting that assembly algorithm is not a useful metric to select software for *de novo* assembly of GBS data.

For the top performing assemblers, CD-HIT and Stacks2, the challenges to obtaining correct assemblies were as expected: allelic polymorphism due to indel variation at a locus likely led to assembly of shorter tracts of true genome fragments into contigs (Stacks2; see Figures S3 S4), and sequence divergence among paralogs and alleles made assemblies appropriately sensitive to the minimum percentage match of sequences within a contig. The choice of minimum match percentage that optimizes over-versus under-assembly will remain a problem for *de novo* assembly until read lengths become much longer than paralogous sequences (allowing them to be placed uniquely in the genome). If not recognized, under- and over-assembly affect downstream analyses, including estimates of population heterozygosity and differentiation (Willis *et al*., 2017; Harvey *et al*., 2015). Of the two, over-assembly is likely preferable for many genomes, as its errors involve closely related, paralogous sequences, which are expected to be rarer than comparable allelic variation at individual loci. Down-stream filtering of loci from population samples may identify likely over-assembled paralogs (O’Leary *et al*., 2018), though limiting over- and under-assembly from the onset is likely desirable. This post-assembly filtering includes excluding loci based on the distribution of read depth across loci and on improbably high heterozygosity given the allele frequencies at a locus (McKinney *et al*., 2017). Ideally, this filtering would be combined with the systematic analysis and comparison of *de novo* assemblies using different percent matches (Willis *et al*., 2017; McCartney-Melstad *et al*., 2019). Tools and methods are available to compare assemblies obtained under different different percent matches (and other settings; Paris *et al*., 2017; Rochette & Catchen, 2017; McCartney-Melstad *et al*., 2019) and these should likely become a standard part of population genomics based on *de novo* assemblies. Additionally, some bioinformatic pipelines provide additional steps to improve the accuracy of assemblies, and may therefore result in more accurate assembled RAD loci than the assembler software would produce on its own (e.g., dDocent; Puritz *et al*., 2014).

Our study indicates CD-HIT is a good choice among currently available programs for *de novo* assembly with varying match percentages, and draws attention to the substantial differences among methods that will be beneficial in evaluating new tools for *de novo* assembly of GBS sequences (e.g., RADProc; Nadukkalam Ravindran *et al*., 2019).

## Data Accessibility

Simulated reads, assembler outputs, and scripts for simulations, assembly, and analysis are available from the Dryad Digital Repository: https://doi.org/10.5061/dryad.8tr03f8.

## Author Contributions

EOA conducted simulations and compiled results. CAB wrote the Perl scripts to compare assemblies to simulated data, conducted the literature review, and performed the CD-HIT assemblies. CH performed the VSEARCH assemblies. ML performed the Stacks, Stacks2, and ABySS assemblies and conducted the literature review. LCM conducted simulations. GR performed Velvet assembles. All authors wrote and reviewed the manuscript.

## Acknowledgements

This project was initiated as part of a computational biology practicum course taught by CAB. All computing was done with the support of the University of Wyoming’s Advanced Research Computing Center, on its IBM System X clusters, Mount Moran (http://n2t.net/ark:/85786/m4159c) and Teton (https://doi.org/10.15786/M2FY47). EOA was supported by the National Science Foundation Graduate Research Fellowship Program and the Wyoming NASA Space Grant Consortium (NASA Grant #NNX15AI08H). MEFL was supported by the University of Wyoming Program in Ecology and her major advisor Holly Ernest’s Wyoming Excellence Chair funds. T. Parchman provided helpful feedback at several stages of the project and provided valuable comments on a draft of the manuscript.

C. Nice also provided valuable comments on a draft of the manuscript.

## Supplementary Material

**Figure S1:**
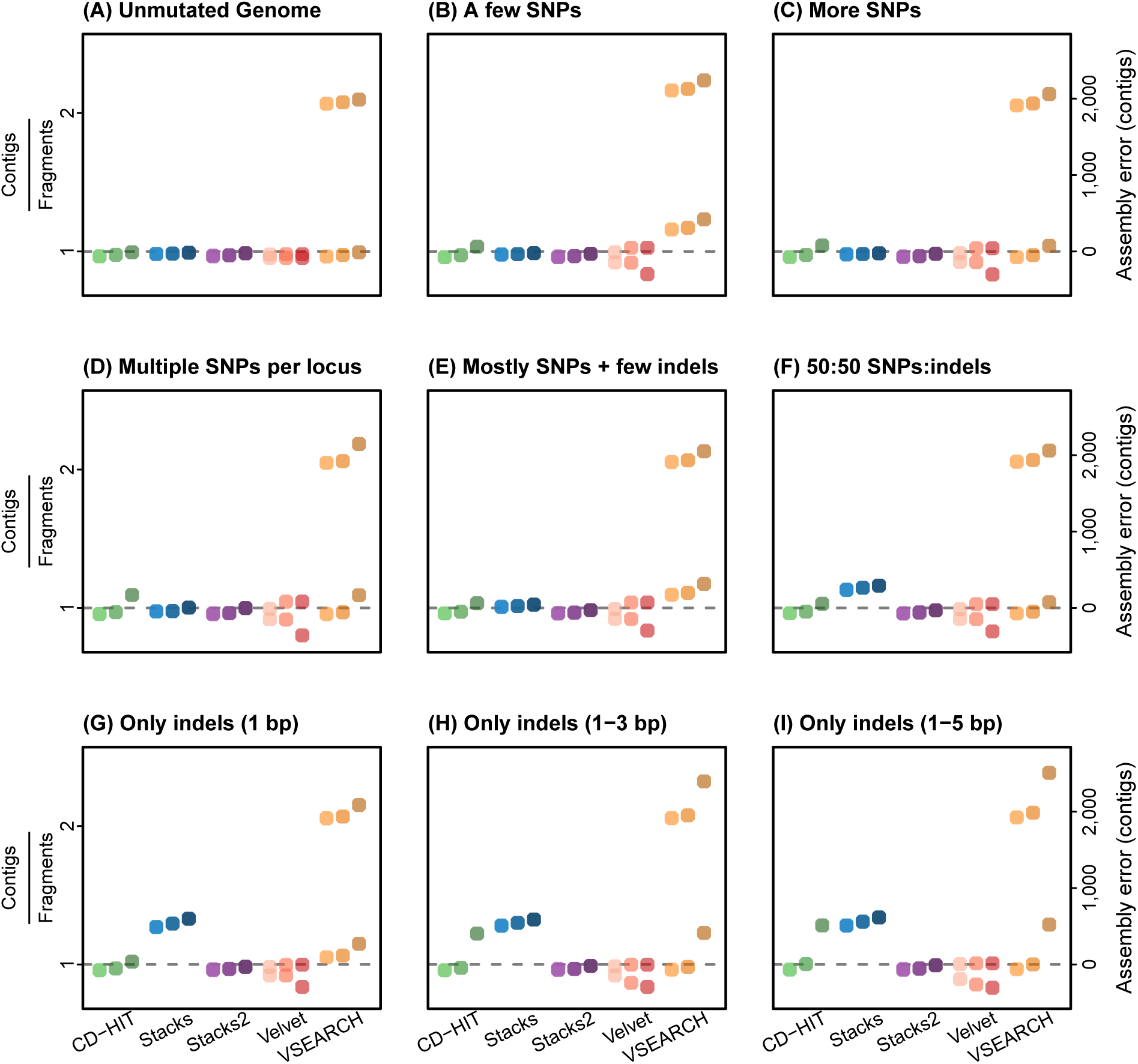
Performance of CD-HIT (green), Stacks (blue), Stacks2 (purple), Velvet (pink), and VSEARCH (orange) on all nine simulations of the *A. thaliana* genome (panes A–I). The ratio of contigs produced by each assembler to the number of unique fragments is used to estimate the degree of under- or over-assembly, with values greater than one representing under-assembly and values less than one representing over-assembly. The second y-axis displays the number of mis-assembled contigs, with positive values representing the number of contigs that are under-assembled and negative values representing the number of contigs that are over-assembled. Perfect assembly, a value of 1, is represented by the gray dashed line. Percent match values used for the assemblies are represented by hue: 90% (light), 94% (medium), and 98% (dark). Assemblers have multiple dots in the same hue when k-mer length affected assembly outcome (see Table S2 for details).

**Figure S2:**
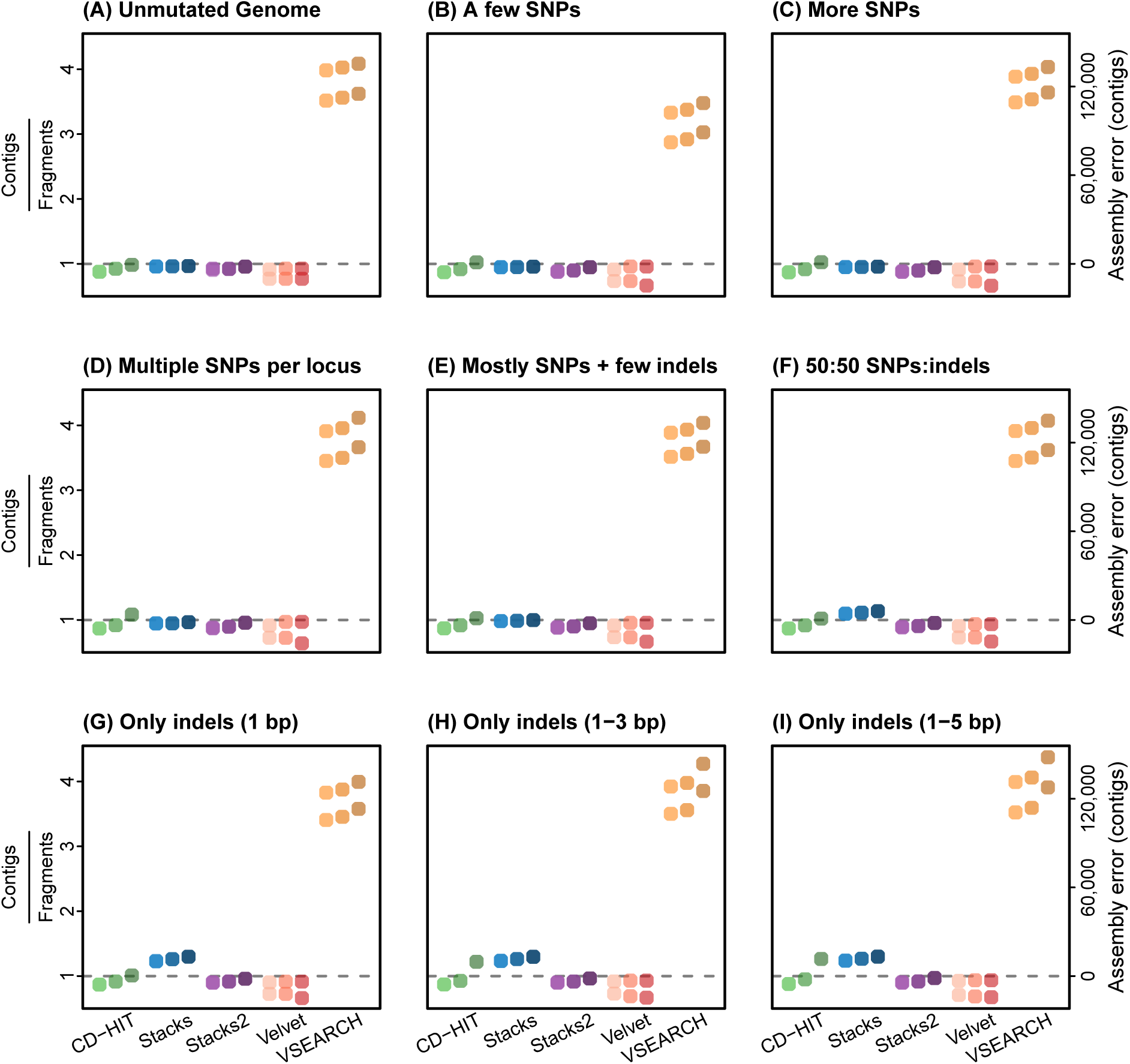
Performance of CD-HIT (green), Stacks (blue), Stacks2 (purple), Velvet (pink), and VSEARCH (orange) on all nine simulations of the *H. sapiens* genome (panes A–I). The ratio of contigs produced by each assembler to the number of unique fragments is used to estimate the degree of under- or over-assembly, with values greater than one representing under-assembly and values less than one representing over-assembly. The second y-axis displays the number of mis-assembled contigs, with positive values representing the number of contigs that are under-assembled and negative values representing the number of contigs that are over-assembled. Perfect assembly, a value of 1, is represented by the gray dashed line. Percent match values used for the assemblies are represented by hue: 90% (light), 94% (medium), and 98% (dark). Assemblers have multiple dots in the same hue when k-mer length affected assembly outcome (see Table S3 for details).

**Figure S3:**
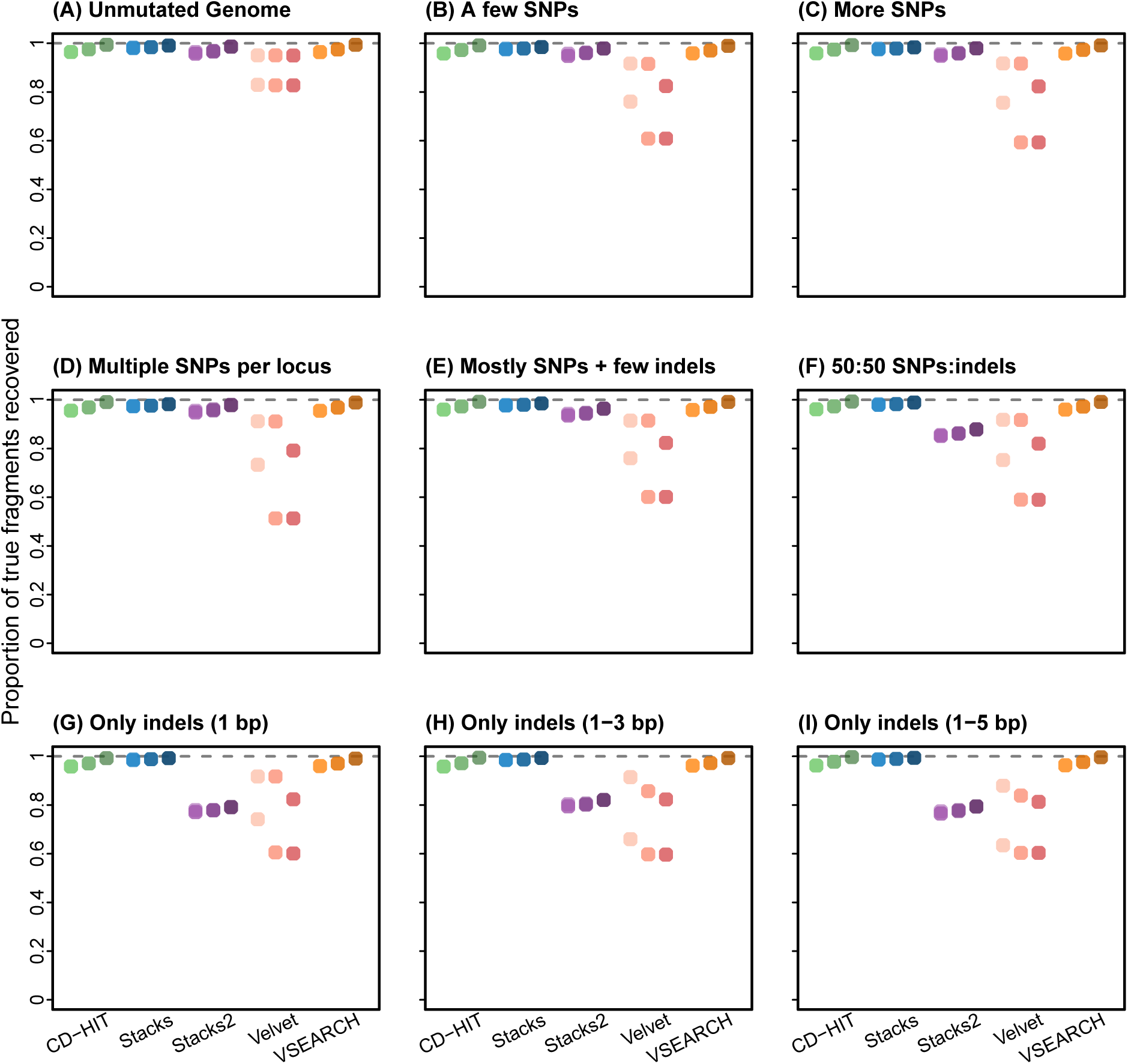
The completeness of assemblies in various simulations (A–I) derived from the *A. thaliana* genome. Completeness was calculated as the proportion of contigs that exactly matched original genome fragments. A value less than 1 indicates that some of the contigs produced were not found in the genome fragments. CD-HIT is green, Stacks is blue, Stacks2 is purple, Velvet is pink, and VSEARCH is orange. The impact of varying the percent match parameter setting is represented by the gradient in the hue of assembler-specific colors (light hue = 90%, medium hue=94% and dark hue=98% match). Assemblers have multiple dots in the same hue when k-mer length affected assembly outcome (see Table S2 for details).

**Figure S4:**
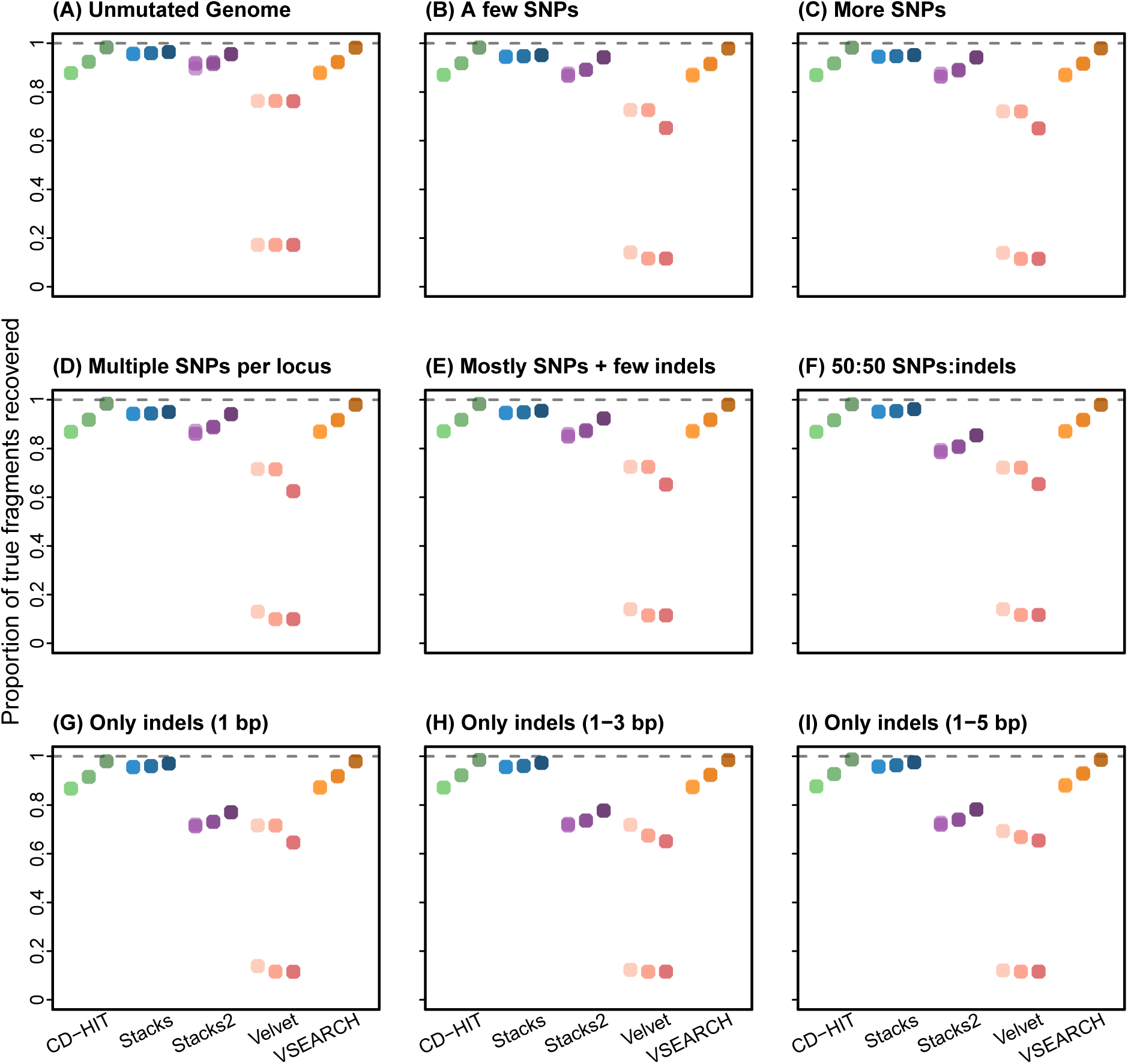
The completeness of assemblies in various simulations (A–I) derived from the *H. sapiens* genome. Completeness was calculated as the proportion of contigs that exactly matched original genome fragments. A value less than 1 indicates that some of the contigs produced were not found in the genome fragments. CD-HIT is green, Stacks is blue, Stacks2 is purple, Velvet is pink, and VSEARCH is orange. The impact of varying the percent match parameter setting is represented by the gradient in the hue of assembler-specific colors (light hue = 90%, medium hue=94% and dark hue=98% match). Assemblers have multiple dots in the same hue when k-mer length affected assembly outcome (see Table S3 for details).

**Table S1:**
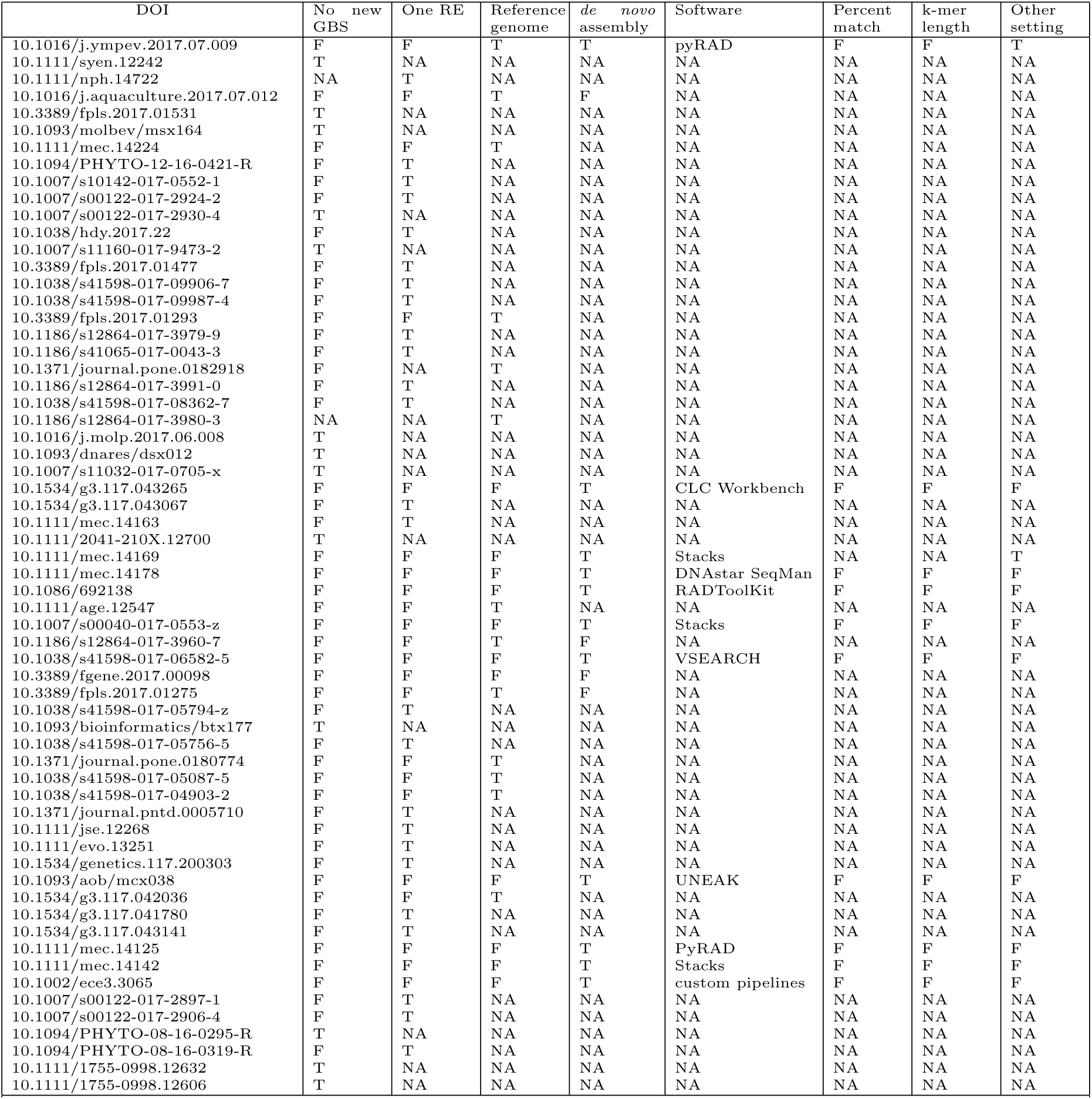
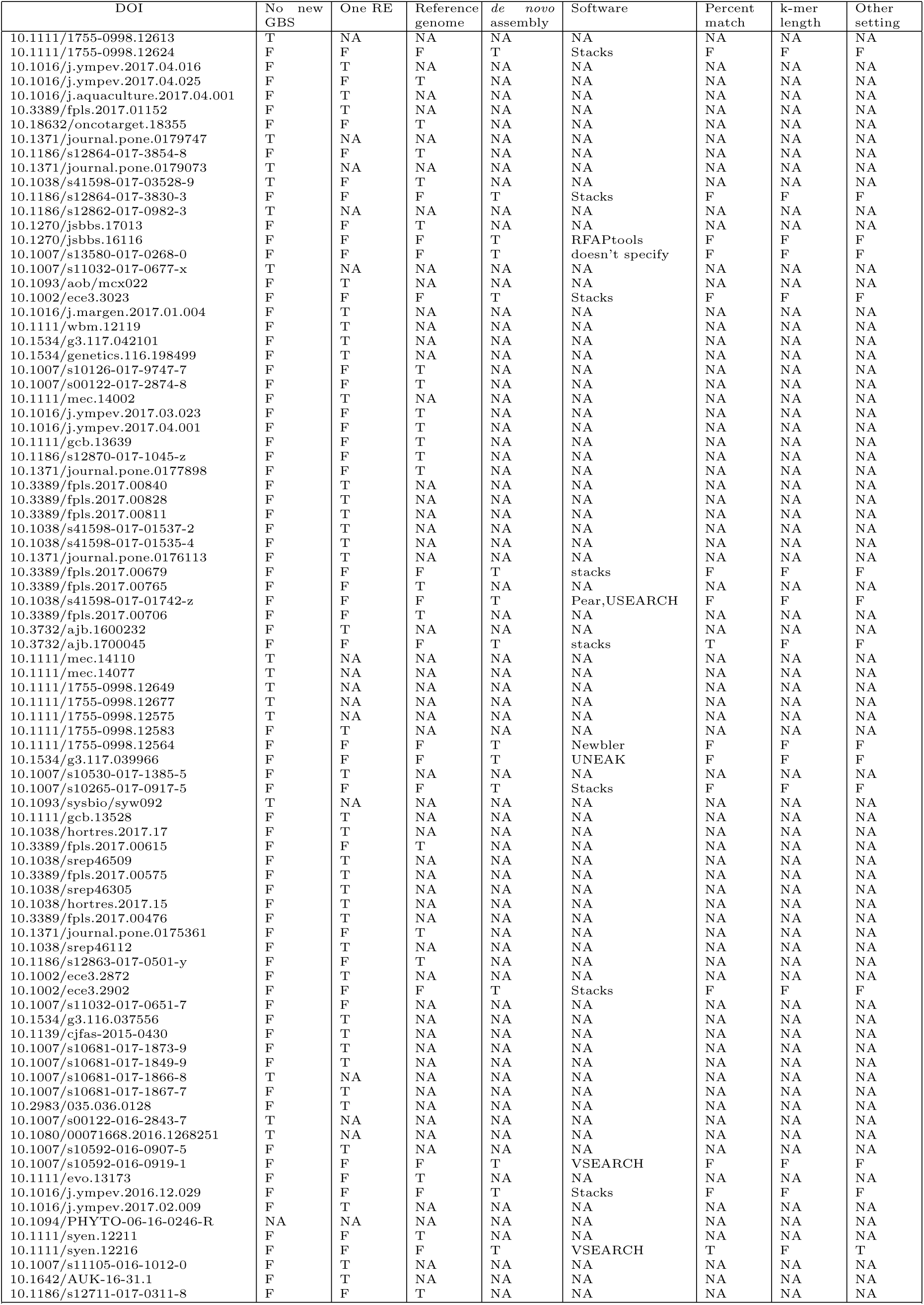
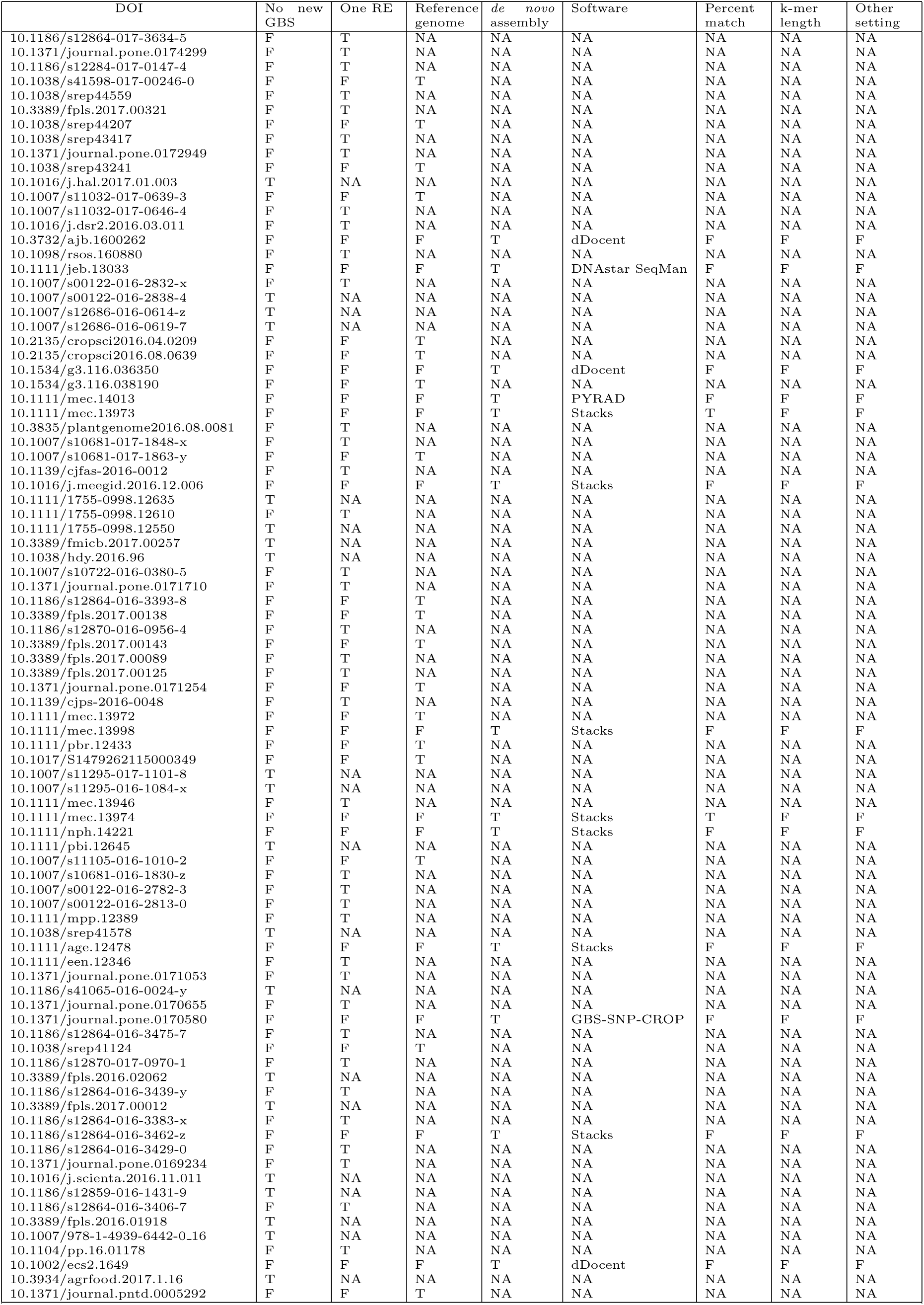
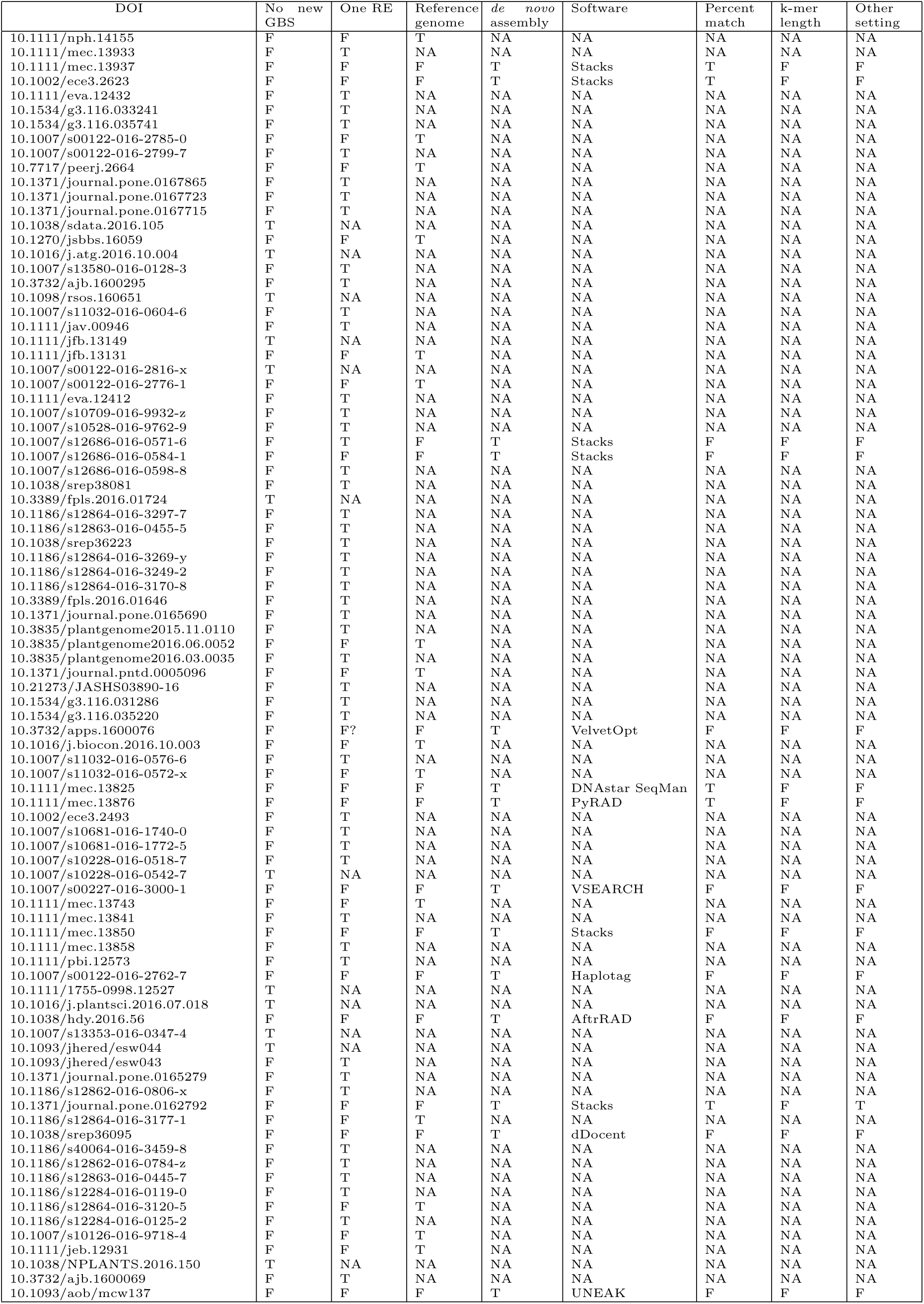
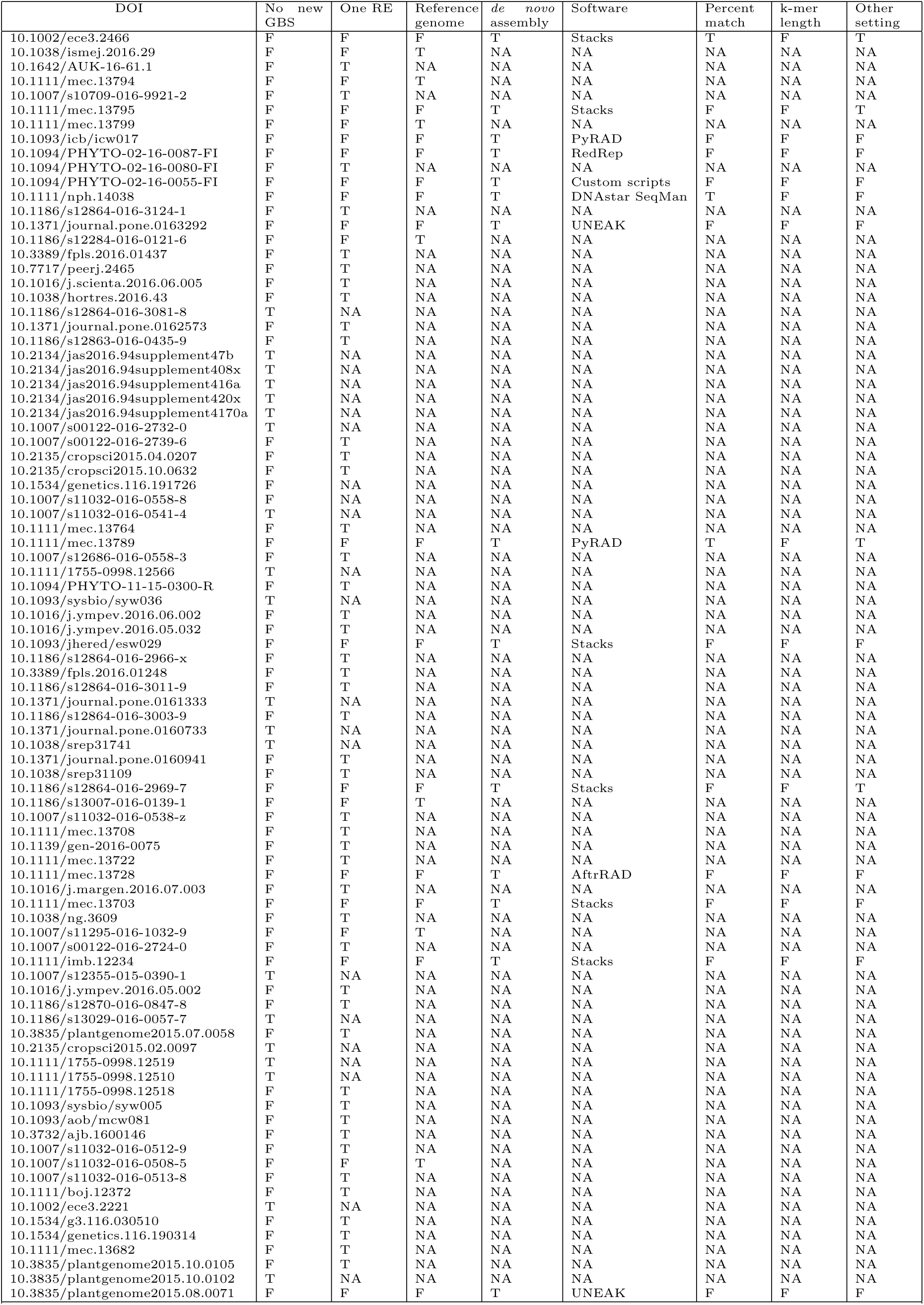
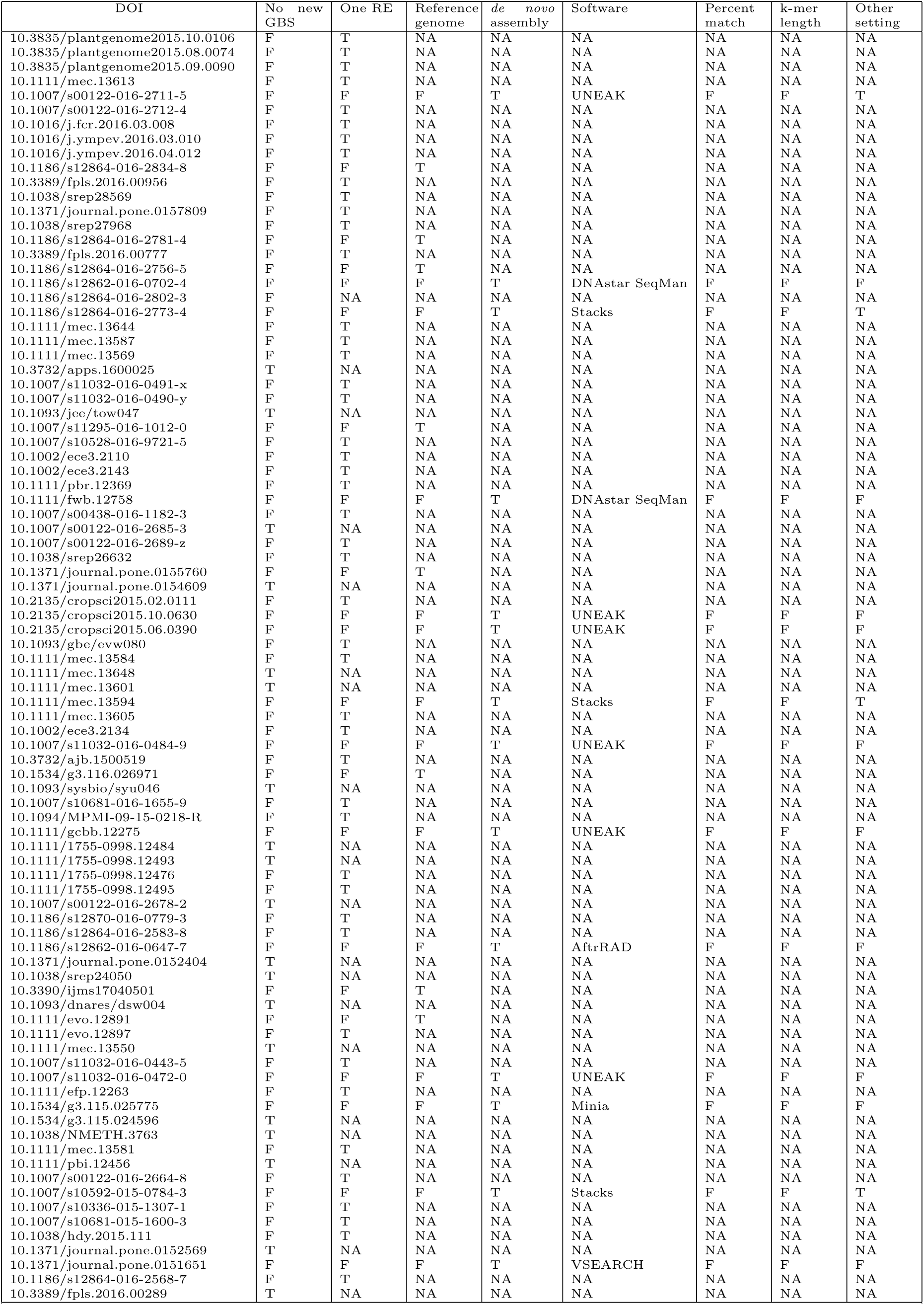
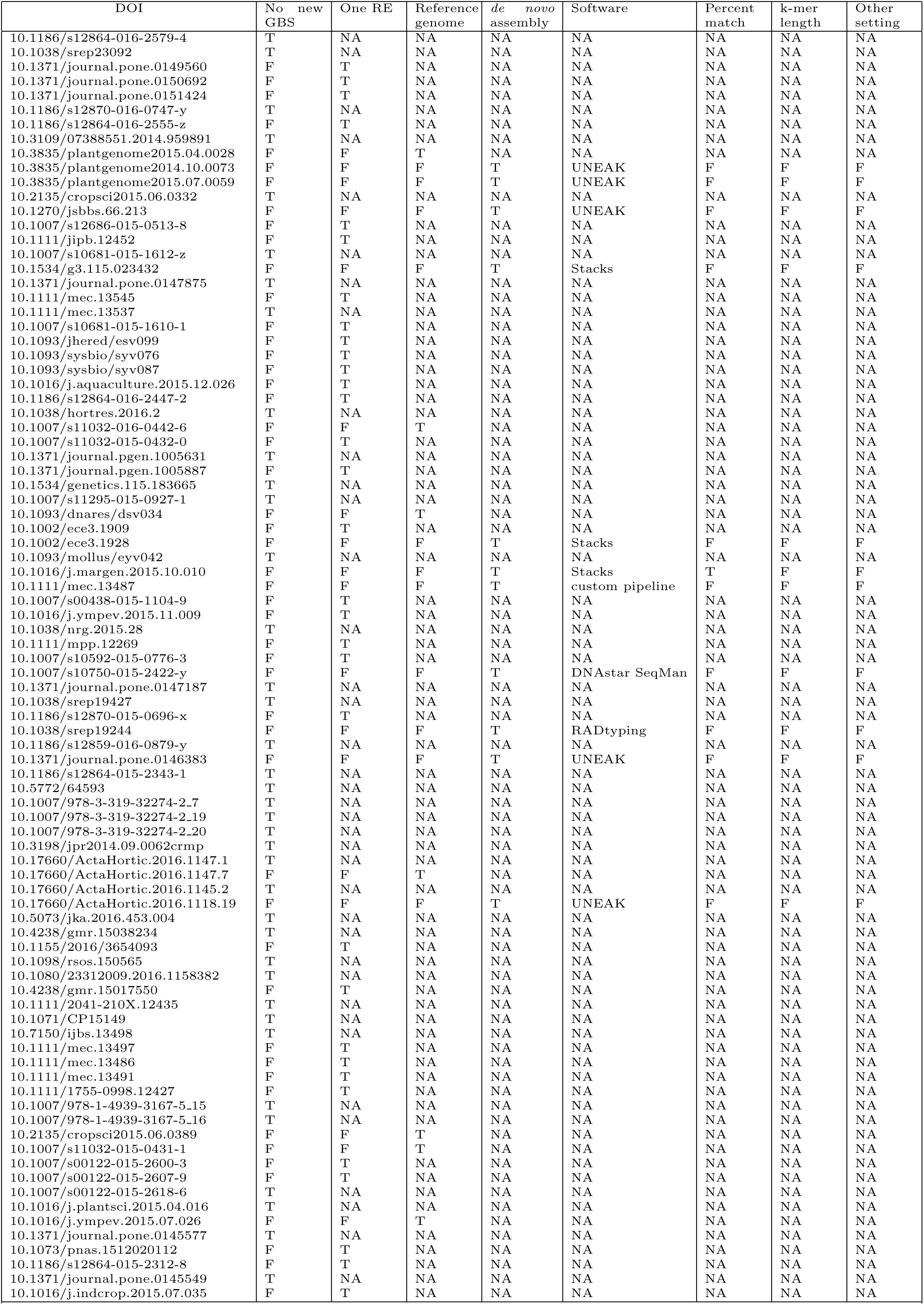
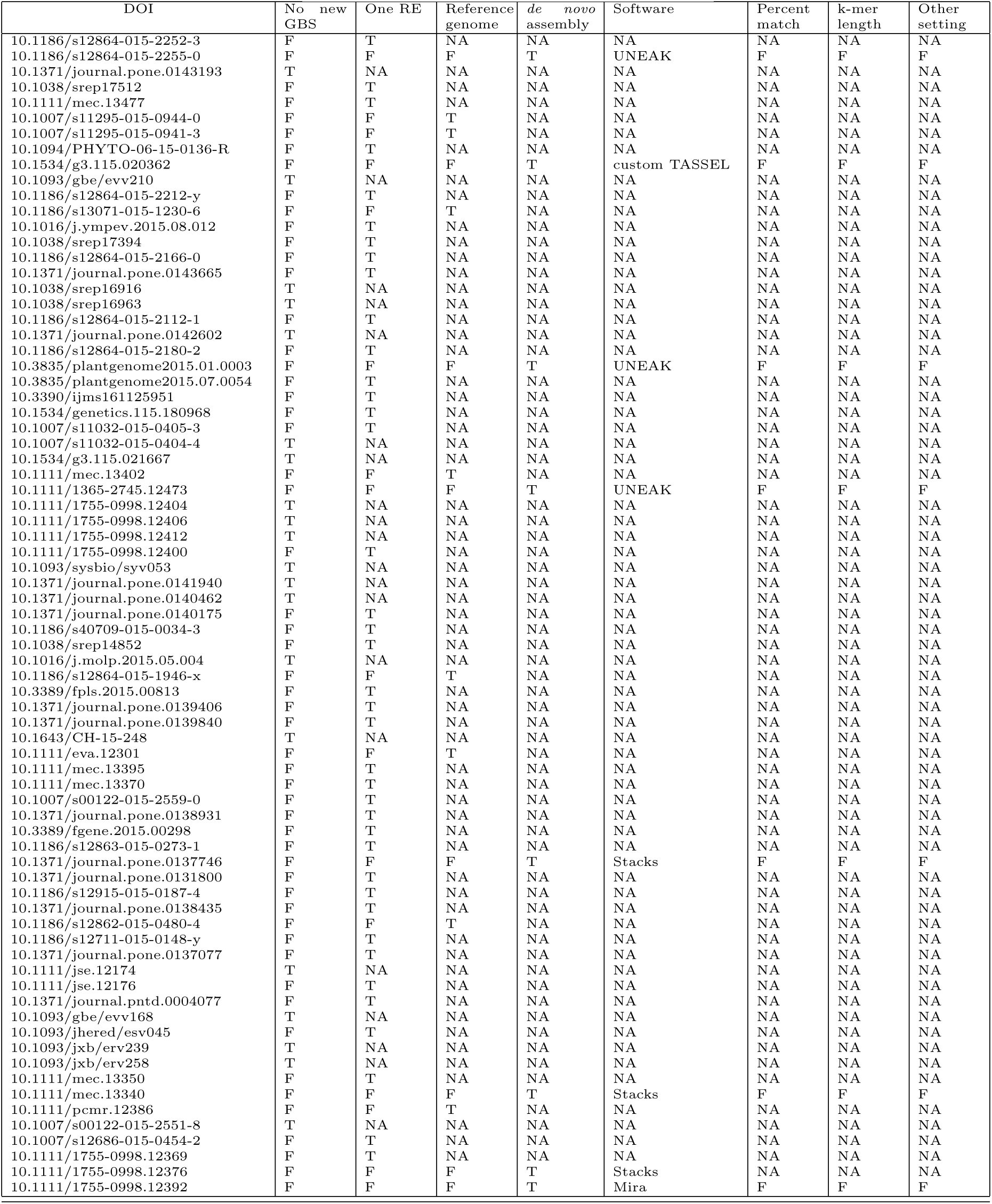
Articles included in our literature review, listed in reverse chronological order. We excluded papers from our literature review that did not present new data, such as review papers (no new GBS), papers that utilized a single restriction enzyme instead of two (one RE), and papers that performed reference-based assembly (reference genome). We sought papers that performed *de novo* assembly (*de novo* assembly), and for these papers we recorded the assembly software used (software), and whether they reported varying parameter settings (percent match, k-mer length, and other setting). For all columns, T = true and F = false. Of the 100 papers meeting our desired criteria, 39 used Stacks (the period we reviewed preceded the release of Stacks2) (Catchen *et al*., 2011), 19 used UNEAK (Lu *et al*., 2013), 11 used VSEARCH (Rognes *et al*., 2016), and 14 used one of the following assemblers: DNASTAR SeqMan (DNAstar, Inc.), dDocent (i.e., CD-HIT) (Puritz *et al*., 2014), or AftrRAD (Sovic *et al*., 2015). The remaining 17 papers each used a unique assembler.

**Table S2:** Assembler results for all simulations derived from the *A. thaliana* genome. Simulations were used to explore the impact of the number of mutations (overall and per locus), the proportion of mutations that were indels and the size of indels, on assembly performance across five different assemblers (i.e., CD-HIT, Stacks, Stacks2, VSEARCH, and Velvet), with varying percent match and k-mer lengths (when appropriate). Detailed descriptions of each simulation can be found in Table 1 of the main text. Assembly results were compared using two different metrics, the proportion of contigs that match genome fragments (completeness) and over-under assembly ratio (contigs:fragments). The completeness proportion includes exact sequence matches as well as close matches (see main text methods for clarification). K-mer length is left blank in the case of CD-HIT entries, because CD-HIT does not use k-mer length as a parameter setting.

| Assembler | Simulation | Percent match | K-mer length | Completeness | Contig/Fragment Ratio |
| --- | --- | --- | --- | --- | --- |
| CD-HIT | Original Genome | 94 | - | 0.98 | 0.98 |
| CD-HIT | Original Genome | 98 | - | 0.99 | 0.99 |
| CD-HIT | Original Genome | 90 | - | 0.96 | 0.96 |
| Stacks | Original Genome | 90 | 15 | 0.98 | 0.98 |
| Stacks | Original Genome | 98 | 15 | 0.99 | 0.99 |
| Stacks | Original Genome | 94 | 15 | 0.98 | 0.98 |
| Stacks | Original Genome | 90 | 31 | 0.98 | 0.98 |
| Stacks | Original Genome | 98 | 31 | 0.99 | 0.99 |
| Stacks | Original Genome | 94 | 31 | 0.98 | 0.98 |
| Stacks | Original Genome | 90 | opt | 0.98 | 0.98 |
| Stacks | Original Genome | 98 | opt | 0.99 | 0.99 |
| Stacks | Original Genome | 94 | opt | 0.98 | 0.98 |
| VSEARCH | Original Genome | 90 | 15 | 0.96 | 2.07 |
| VSEARCH | Original Genome | 90 | 8 | 0.96 | 0.96 |
| VSEARCH | Original Genome | 94 | 15 | 0.97 | 2.08 |
| VSEARCH | Original Genome | 94 | 8 | 0.97 | 0.97 |
| VSEARCH | Original Genome | 98 | 15 | 0.99 | 2.10 |
| VSEARCH | Original Genome | 98 | 8 | 0.99 | 0.99 |
| Velvet | Original Genome | 98 | 15 | 0.83 | 0.98 |
| Velvet | Original Genome | 94 | 15 | 0.83 | 0.98 |
| Velvet | Original Genome | 90 | 15 | 0.83 | 0.98 |
| Velvet | Original Genome | 98 | 31 | 0.95 | 0.95 |
| Velvet | Original Genome | 94 | 31 | 0.95 | 0.95 |
| Velvet | Original Genome | 90 | 31 | 0.95 | 0.95 |
| Stacks2 | Original Genome | 98 | opt | 0.99 | 0.99 |
| Stacks2 | Original Genome | 90 | 15 | 0.95 | 0.96 |
| Stacks2 | Original Genome | 94 | 15 | 0.96 | 0.97 |
| Stacks2 | Original Genome | 94 | 31 | 0.97 | 0.97 |
| Stacks2 | Original Genome | 98 | 31 | 0.99 | 0.99 |
| Stacks2 | Original Genome | 94 | opt | 0.96 | 0.97 |
| Stacks2 | Original Genome | 90 | opt | 0.95 | 0.96 |
| Stacks2 | Original Genome | 90 | 31 | 0.96 | 0.97 |
| Stacks2 | Original Genome | 98 | 15 | 0.98 | 0.99 |
| CD-HIT | A few SNPs | 94 | - | 0.97 | 0.97 |
| CD-HIT | A few SNPs | 98 | - | 0.99 | 1.03 |

Table S2 – continued from previous page
| Assembler | Simulation | Percent match | K-mer length | Completeness | Contig/Fragment Ratio |
| --- | --- | --- | --- | --- | --- |
| CD-HIT | A few SNPs | 90 | - | 0.96 | 0.96 |
| Stacks | A few SNPs | 90 | 15 | 0.98 | 0.98 |
| Stacks | A few SNPs | 98 | 15 | 0.98 | 0.99 |
| Stacks | A few SNPs | 94 | 15 | 0.98 | 0.98 |
| Stacks | A few SNPs | 90 | 31 | 0.98 | 0.98 |
| Stacks | A few SNPs | 98 | 31 | 0.98 | 0.99 |
| Stacks | A few SNPs | 94 | 31 | 0.98 | 0.98 |
| Stacks | A few SNPs | 90 | opt | 0.98 | 0.98 |
| Stacks | A few SNPs | 98 | opt | 0.98 | 0.99 |
| Stacks | A few SNPs | 94 | opt | 0.98 | 0.98 |
| VSEARCH | A few SNPs | 90 | 15 | 0.96 | 2.16 |
| VSEARCH | A few SNPs | 90 | 8 | 0.96 | 1.16 |
| VSEARCH | A few SNPs | 94 | 15 | 0.97 | 2.17 |
| VSEARCH | A few SNPs | 94 | 8 | 0.97 | 1.17 |
| VSEARCH | A few SNPs | 98 | 15 | 0.99 | 2.24 |
| VSEARCH | A few SNPs | 98 | 8 | 0.99 | 1.23 |
| Velvet | A few SNPs | 98 | 15 | 0.61 | 1.03 |
| Velvet | A few SNPs | 94 | 15 | 0.61 | 1.03 |
| Velvet | A few SNPs | 90 | 15 | 0.76 | 0.99 |
| Velvet | A few SNPs | 98 | 31 | 0.82 | 0.84 |
| Velvet | A few SNPs | 94 | 31 | 0.92 | 0.92 |
| Velvet | A few SNPs | 90 | 31 | 0.92 | 0.92 |
| Stacks2 | A few SNPs | 94 | 15 | 0.96 | 0.96 |
| Stacks2 | A few SNPs | 90 | 31 | 0.96 | 0.96 |
| Stacks2 | A few SNPs | 98 | 15 | 0.98 | 0.98 |
| Stacks2 | A few SNPs | 94 | 31 | 0.96 | 0.97 |
| Stacks2 | A few SNPs | 98 | 31 | 0.98 | 0.98 |
| Stacks2 | A few SNPs | 98 | opt | 0.98 | 0.98 |
| Stacks2 | A few SNPs | 94 | opt | 0.96 | 0.97 |
| Stacks2 | A few SNPs | 90 | opt | 0.95 | 0.96 |
| Stacks2 | A few SNPs | 90 | 15 | 0.95 | 0.96 |
| CD-HIT | More SNPs | 94 | - | 0.97 | 0.97 |
| CD-HIT | More SNPs | 98 | - | 0.99 | 1.04 |
| CD-HIT | More SNPs | 90 | - | 0.96 | 0.96 |
| Stacks | More SNPs | 90 | 15 | 0.98 | 0.98 |
| Stacks | More SNPs | 98 | 15 | 0.98 | 0.99 |
| Stacks | More SNPs | 94 | 15 | 0.98 | 0.98 |
| Stacks | More SNPs | 90 | 31 | 0.98 | 0.98 |
| Stacks | More SNPs | 98 | 31 | 0.98 | 0.99 |
| Stacks | More SNPs | 94 | 31 | 0.98 | 0.98 |
| Stacks | More SNPs | 90 | opt | 0.98 | 0.98 |
| Stacks | More SNPs | 98 | opt | 0.98 | 0.99 |
| Stacks | More SNPs | 94 | opt | 0.98 | 0.98 |
| VSEARCH | More SNPs | 90 | 15 | 0.96 | 2.05 |
| VSEARCH | More SNPs | 90 | 8 | 0.96 | 0.96 |
| VSEARCH | More SNPs | 94 | 15 | 0.97 | 2.07 |
| VSEARCH | More SNPs | 94 | 8 | 0.97 | 0.97 |
| VSEARCH | More SNPs | 98 | 15 | 0.99 | 2.14 |
| VSEARCH | More SNPs | 98 | 8 | 0.99 | 1.04 |
| Velvet | More SNPs | 98 | 15 | 0.59 | 1.02 |

Table S2 – continued from previous page
| Assembler | Simulation | Percent match | K-mer length | Completeness | Contig/Fragment Ratio |
| --- | --- | --- | --- | --- | --- |
| Velvet | More SNPs | 94 | 15 | 0.59 | 1.02 |
| Velvet | More SNPs | 90 | 15 | 0.76 | 0.99 |
| Velvet | More SNPs | 98 | 31 | 0.82 | 0.83 |
| Velvet | More SNPs | 94 | 31 | 0.92 | 0.92 |
| Velvet | More SNPs | 90 | 31 | 0.92 | 0.92 |
| Stacks2 | More SNPs | 98 | opt | 0.98 | 0.98 |
| Stacks2 | More SNPs | 90 | 31 | 0.96 | 0.96 |
| Stacks2 | More SNPs | 98 | 15 | 0.98 | 0.98 |
| Stacks2 | More SNPs | 90 | opt | 0.95 | 0.96 |
| Stacks2 | More SNPs | 94 | 15 | 0.96 | 0.96 |
| Stacks2 | More SNPs | 94 | 31 | 0.96 | 0.97 |
| Stacks2 | More SNPs | 94 | opt | 0.96 | 0.96 |
| Stacks2 | More SNPs | 90 | 15 | 0.95 | 0.96 |
| Stacks2 | More SNPs | 98 | 31 | 0.98 | 0.98 |
| CD-HIT | Multiple SNPs per locus | 94 | - | 0.97 | 0.97 |
| CD-HIT | Multiple SNPs per locus | 98 | - | 0.99 | 1.09 |
| CD-HIT | Multiple SNPs per locus | 90 | - | 0.96 | 0.96 |
| Stacks | Multiple SNPs per locus | 90 | 15 | 0.97 | 0.97 |
| Stacks | Multiple SNPs per locus | 98 | 15 | 0.98 | 1.00 |
| Stacks | Multiple SNPs per locus | 94 | 15 | 0.98 | 0.98 |
| Stacks | Multiple SNPs per locus | 90 | 31 | 0.97 | 0.98 |
| Stacks | Multiple SNPs per locus | 98 | 31 | 0.98 | 1.00 |
| Stacks | Multiple SNPs per locus | 94 | 31 | 0.98 | 0.98 |
| Stacks | Multiple SNPs per locus | 90 | opt | 0.97 | 0.97 |
| Stacks | Multiple SNPs per locus | 98 | opt | 0.98 | 1.00 |
| Stacks | Multiple SNPs per locus | 94 | opt | 0.98 | 0.98 |
| VSEARCH | Multiple SNPs per locus | 90 | 15 | 0.96 | 2.05 |
| VSEARCH | Multiple SNPs per locus | 90 | 8 | 0.96 | 0.96 |
| VSEARCH | Multiple SNPs per locus | 94 | 15 | 0.97 | 2.06 |
| VSEARCH | Multiple SNPs per locus | 94 | 8 | 0.97 | 0.97 |
| VSEARCH | Multiple SNPs per locus | 98 | 15 | 0.99 | 2.18 |
| VSEARCH | Multiple SNPs per locus | 98 | 8 | 0.99 | 1.09 |
| Velvet | Multiple SNPs per locus | 98 | 15 | 0.51 | 1.05 |
| Velvet | Multiple SNPs per locus | 94 | 15 | 0.51 | 1.05 |
| Velvet | Multiple SNPs per locus | 90 | 15 | 0.73 | 0.99 |
| Velvet | Multiple SNPs per locus | 98 | 31 | 0.79 | 0.80 |
| Velvet | Multiple SNPs per locus | 94 | 31 | 0.91 | 0.92 |
| Velvet | Multiple SNPs per locus | 90 | 31 | 0.91 | 0.92 |
| Stacks2 | Multiple SNPs per locus | 98 | 15 | 0.98 | 1.00 |
| Stacks2 | Multiple SNPs per locus | 94 | opt | 0.95 | 0.96 |
| Stacks2 | Multiple SNPs per locus | 90 | 31 | 0.95 | 0.96 |
| Stacks2 | Multiple SNPs per locus | 90 | 15 | 0.95 | 0.96 |
| Stacks2 | Multiple SNPs per locus | 98 | 31 | 0.98 | 1.00 |
| Stacks2 | Multiple SNPs per locus | 94 | 31 | 0.96 | 0.96 |
| Stacks2 | Multiple SNPs per locus | 90 | opt | 0.94 | 0.95 |
| Stacks2 | Multiple SNPs per locus | 94 | 15 | 0.96 | 0.96 |
| Stacks2 | Multiple SNPs per locus | 98 | opt | 0.98 | 1.00 |
| CD-HIT | Mostly SNPs + few indels | 94 | - | 0.97 | 0.97 |
| CD-HIT | Mostly SNPs + few indels | 98 | - | 0.99 | 1.03 |
| CD-HIT | Mostly SNPs + few indels | 90 | - | 0.96 | 0.96 |

Table S2 – continued from previous page
| Assembler | Simulation | Percent match | K-mer length | Completeness | Contig/Fragment Ratio |
| --- | --- | --- | --- | --- | --- |
| Stacks | Mostly SNPs + few indels | 90 | 15 | 0.98 | 1.01 |
| Stacks | Mostly SNPs + few indels | 98 | 15 | 0.99 | 1.02 |
| Stacks | Mostly SNPs + few indels | 94 | 15 | 0.98 | 1.01 |
| Stacks | Mostly SNPs + few indels | 90 | 31 | 0.98 | 1.01 |
| Stacks | Mostly SNPs + few indels | 98 | 31 | 0.99 | 1.02 |
| Stacks | Mostly SNPs + few indels | 94 | 31 | 0.98 | 1.01 |
| Stacks | Mostly SNPs + few indels | 90 | opt | 0.98 | 1.01 |
| Stacks | Mostly SNPs + few indels | 98 | opt | 0.99 | 1.02 |
| Stacks | Mostly SNPs + few indels | 94 | opt | 0.98 | 1.01 |
| VSEARCH | Mostly SNPs + few indels | 90 | 15 | 0.96 | 2.05 |
| VSEARCH | Mostly SNPs + few indels | 90 | 8 | 0.96 | 1.10 |
| VSEARCH | Mostly SNPs + few indels | 94 | 15 | 0.97 | 2.07 |
| VSEARCH | Mostly SNPs + few indels | 94 | 8 | 0.97 | 1.11 |
| VSEARCH | Mostly SNPs + few indels | 98 | 15 | 0.99 | 2.13 |
| VSEARCH | Mostly SNPs + few indels | 98 | 8 | 0.99 | 1.17 |
| Velvet | Mostly SNPs + few indels | 98 | 15 | 0.60 | 1.04 |
| Velvet | Mostly SNPs + few indels | 94 | 15 | 0.60 | 1.04 |
| Velvet | Mostly SNPs + few indels | 90 | 15 | 0.75 | 0.99 |
| Velvet | Mostly SNPs + few indels | 98 | 31 | 0.81 | 0.84 |
| Velvet | Mostly SNPs + few indels | 94 | 31 | 0.90 | 0.92 |
| Velvet | Mostly SNPs + few indels | 90 | 31 | 0.90 | 0.92 |
| Stacks2 | Mostly SNPs + few indels | 94 | 31 | 0.93 | 0.97 |
| Stacks2 | Mostly SNPs + few indels | 90 | opt | 0.92 | 0.96 |
| Stacks2 | Mostly SNPs + few indels | 90 | 31 | 0.92 | 0.96 |
| Stacks2 | Mostly SNPs + few indels | 98 | opt | 0.94 | 0.98 |
| Stacks2 | Mostly SNPs + few indels | 98 | 15 | 0.94 | 0.98 |
| Stacks2 | Mostly SNPs + few indels | 90 | 15 | 0.92 | 0.96 |
| Stacks2 | Mostly SNPs + few indels | 98 | 31 | 0.94 | 0.98 |
| Stacks2 | Mostly SNPs + few indels | 94 | opt | 0.92 | 0.96 |
| Stacks2 | Mostly SNPs + few indels | 94 | 15 | 0.92 | 0.97 |
| CD-HIT | 50:50 SNPs:indels | 94 | - | 0.97 | 0.97 |
| CD-HIT | 50:50 SNPs:indels | 98 | - | 0.99 | 1.03 |
| CD-HIT | 50:50 SNPs:indels | 90 | - | 0.96 | 0.96 |
| Stacks | 50:50 SNPs:indels | 90 | 15 | 0.98 | 1.13 |
| Stacks | 50:50 SNPs:indels | 98 | 15 | 0.99 | 1.16 |
| Stacks | 50:50 SNPs:indels | 94 | 15 | 0.98 | 1.15 |
| Stacks | 50:50 SNPs:indels | 90 | 31 | 0.98 | 1.13 |
| Stacks | 50:50 SNPs:indels | 98 | 31 | 0.99 | 1.16 |
| Stacks | 50:50 SNPs:indels | 94 | 31 | 0.98 | 1.15 |
| Stacks | 50:50 SNPs:indels | 90 | opt | 0.98 | 1.13 |
| Stacks | 50:50 SNPs:indels | 98 | opt | 0.99 | 1.16 |
| Stacks | 50:50 SNPs:indels | 94 | opt | 0.98 | 1.15 |
| VSEARCH | 50:50 SNPs:indels | 90 | 15 | 0.96 | 2.06 |
| VSEARCH | 50:50 SNPs:indels | 90 | 8 | 0.96 | 0.96 |
| VSEARCH | 50:50 SNPs:indels | 94 | 15 | 0.97 | 2.07 |
| VSEARCH | 50:50 SNPs:indels | 94 | 8 | 0.97 | 0.97 |
| VSEARCH | 50:50 SNPs:indels | 98 | 15 | 0.99 | 2.14 |
| VSEARCH | 50:50 SNPs:indels | 98 | 8 | 0.99 | 1.04 |
| Velvet | 50:50 SNPs:indels | 98 | 15 | 0.58 | 1.03 |
| Velvet | 50:50 SNPs:indels | 94 | 15 | 0.58 | 1.03 |

Table S2 – continued from previous page
| Assembler | Simulation | Percent match | K-mer length | Completeness | Contig/Fragment Ratio |
| --- | --- | --- | --- | --- | --- |
| Velvet | 50:50 SNPs:indels | 90 | 15 | 0.71 | 0.99 |
| Velvet | 50:50 SNPs:indels | 98 | 31 | 0.80 | 0.83 |
| Velvet | 50:50 SNPs:indels | 94 | 31 | 0.88 | 0.92 |
| Velvet | 50:50 SNPs:indels | 90 | 31 | 0.88 | 0.92 |
| Stacks2 | 50:50 SNPs:indels | 90 | 31 | 0.79 | 0.96 |
| Stacks2 | 50:50 SNPs:indels | 98 | opt | 0.81 | 0.98 |
| Stacks2 | 50:50 SNPs:indels | 94 | 31 | 0.79 | 0.97 |
| Stacks2 | 50:50 SNPs:indels | 94 | opt | 0.79 | 0.97 |
| Stacks2 | 50:50 SNPs:indels | 94 | 15 | 0.79 | 0.97 |
| Stacks2 | 50:50 SNPs:indels | 98 | 15 | 0.81 | 0.98 |
| Stacks2 | 50:50 SNPs:indels | 90 | opt | 0.78 | 0.96 |
| Stacks2 | 50:50 SNPs:indels | 98 | 31 | 0.81 | 0.98 |
| Stacks2 | 50:50 SNPs:indels | 90 | 15 | 0.78 | 0.96 |
| CD-HIT | Only indels | 94 | - | 0.97 | 0.97 |
| CD-HIT | Only indels | 98 | - | 0.99 | 1.02 |
| CD-HIT | Only indels | 90 | - | 0.96 | 0.96 |
| Stacks | Only indels | 90 | 15 | 0.98 | 1.27 |
| Stacks | Only indels | 98 | 15 | 0.99 | 1.33 |
| Stacks | Only indels | 94 | 15 | 0.99 | 1.30 |
| Stacks | Only indels | 90 | 31 | 0.99 | 1.27 |
| Stacks | Only indels | 98 | 31 | 0.99 | 1.33 |
| Stacks | Only indels | 94 | 31 | 0.99 | 1.30 |
| Stacks | Only indels | 90 | opt | 0.98 | 1.27 |
| Stacks | Only indels | 98 | opt | 0.99 | 1.33 |
| Stacks | Only indels | 94 | opt | 0.99 | 1.30 |
| VSEARCH | Only indels | 90 | 8 | 0.96 | 1.05 |
| VSEARCH | Only indels | 94 | 15 | 0.97 | 2.07 |
| VSEARCH | Only indels | 94 | 8 | 0.97 | 1.06 |
| VSEARCH | Only indels | 98 | 15 | 0.99 | 2.15 |
| VSEARCH | Only indels | 98 | 8 | 0.99 | 1.15 |
| VSEARCH | Only indels | 90 | 15 | 0.96 | 2.06 |
| Velvet | Only indels | 98 | 15 | 0.59 | 1.00 |
| Velvet | Only indels | 94 | 15 | 0.59 | 1.00 |
| Velvet | Only indels | 90 | 15 | 0.67 | 0.98 |
| Velvet | Only indels | 98 | 31 | 0.78 | 0.84 |
| Velvet | Only indels | 94 | 31 | 0.83 | 0.92 |
| Velvet | Only indels | 90 | 31 | 0.83 | 0.92 |
| Stacks2 | Only indels | 94 | opt | 0.63 | 0.97 |
| Stacks2 | Only indels | 98 | 31 | 0.64 | 0.98 |
| Stacks2 | Only indels | 90 | 15 | 0.62 | 0.96 |
| Stacks2 | Only indels | 90 | 31 | 0.63 | 0.97 |
| Stacks2 | Only indels | 90 | opt | 0.62 | 0.96 |
| Stacks2 | Only indels | 94 | 31 | 0.63 | 0.97 |
| Stacks2 | Only indels | 98 | 15 | 0.64 | 0.98 |
| Stacks2 | Only indels | 98 | opt | 0.64 | 0.98 |
| Stacks2 | Only indels | 94 | 15 | 0.63 | 0.97 |
| CD-HIT | Only indels, short-mid length | 94 | - | 0.97 | 0.98 |
| CD-HIT | Only indels, short-mid length | 98 | - | 0.99 | 1.22 |
| CD-HIT | Only indels, short-mid length | 90 | - | 0.96 | 0.96 |
| Stacks | Only indels, short-mid length | 90 | 15 | 0.98 | 1.28 |

Table S2 – continued from previous page
| Assembler | Simulation | Percent match | K-mer length | Completeness | Contig/Fragment Ratio |
| --- | --- | --- | --- | --- | --- |
| Stacks | Only indels, short-mid length | 98 | 15 | 0.99 | 1.32 |
| Stacks | Only indels, short-mid length | 94 | 15 | 0.99 | 1.30 |
| Stacks | Only indels, short-mid length | 90 | 31 | 0.99 | 1.28 |
| Stacks | Only indels, short-mid length | 98 | 31 | 0.99 | 1.32 |
| Stacks | Only indels, short-mid length | 94 | 31 | 0.99 | 1.30 |
| Stacks | Only indels, short-mid length | 90 | opt | 0.98 | 1.28 |
| Stacks | Only indels, short-mid length | 98 | opt | 0.99 | 1.32 |
| Stacks | Only indels, short-mid length | 94 | opt | 0.99 | 1.30 |
| VSEARCH | Only indels, short-mid length | 90 | 8 | 0.96 | 0.96 |
| VSEARCH | Only indels, short-mid length | 94 | 15 | 0.97 | 2.08 |
| VSEARCH | Only indels, short-mid length | 94 | 8 | 0.97 | 0.98 |
| VSEARCH | Only indels, short-mid length | 98 | 15 | 0.99 | 2.32 |
| VSEARCH | Only indels, short-mid length | 98 | 8 | 0.99 | 1.23 |
| VSEARCH | Only indels, short-mid length | 90 | 15 | 0.96 | 2.06 |
| Velvet | Only indels, short-mid length | 98 | 15 | 0.58 | 1.00 |
| Velvet | Only indels, short-mid length | 94 | 15 | 0.58 | 1.00 |
| Velvet | Only indels, short-mid length | 90 | 15 | 0.62 | 0.99 |
| Velvet | Only indels, short-mid length | 98 | 31 | 0.77 | 0.84 |
| Velvet | Only indels, short-mid length | 94 | 31 | 0.79 | 0.87 |
| Velvet | Only indels, short-mid length | 90 | 31 | 0.82 | 0.92 |
| Stacks2 | Only indels, short-mid length | 94 | 15 | 0.64 | 0.97 |
| Stacks2 | Only indels, short-mid length | 98 | 15 | 0.65 | 0.99 |
| Stacks2 | Only indels, short-mid length | 90 | 31 | 0.64 | 0.97 |
| Stacks2 | Only indels, short-mid length | 90 | opt | 0.63 | 0.96 |
| Stacks2 | Only indels, short-mid length | 94 | 31 | 0.64 | 0.97 |
| Stacks2 | Only indels, short-mid length | 90 | 15 | 0.63 | 0.96 |
| Stacks2 | Only indels, short-mid length | 98 | opt | 0.65 | 0.99 |
| Stacks2 | Only indels, short-mid length | 94 | opt | 0.64 | 0.97 |
| Stacks2 | Only indels, short-mid length | 98 | 31 | 0.65 | 0.99 |
| CD-HIT | Only indels, short-long length | 94 | - | 0.98 | 1.00 |
| CD-HIT | Only indels, short-long length | 98 | - | 1.00 | 1.28 |
| CD-HIT | Only indels, short-long length | 90 | - | 0.96 | 0.96 |
| Stacks | Only indels, short-long length | 90 | 15 | 0.99 | 1.28 |
| Stacks | Only indels, short-long length | 98 | 15 | 0.99 | 1.34 |
| Stacks | Only indels, short-long length | 94 | 15 | 0.99 | 1.31 |
| Stacks | Only indels, short-long length | 90 | 31 | 0.99 | 1.28 |
| Stacks | Only indels, short-long length | 98 | 31 | 0.99 | 1.34 |
| Stacks | Only indels, short-long length | 94 | 31 | 0.99 | 1.31 |
| Stacks | Only indels, short-long length | 90 | opt | 0.99 | 1.28 |
| Stacks | Only indels, short-long length | 98 | opt | 0.99 | 1.34 |
| Stacks | Only indels, short-long length | 94 | opt | 0.99 | 1.31 |
| VSEARCH | Only indels, short-long length | 90 | 15 | 0.96 | 2.06 |
| VSEARCH | Only indels, short-long length | 90 | 8 | 0.96 | 0.97 |
| VSEARCH | Only indels, short-long length | 94 | 15 | 0.98 | 2.10 |
| VSEARCH | Only indels, short-long length | 94 | 8 | 0.98 | 1.00 |
| VSEARCH | Only indels, short-long length | 98 | 15 | 1.00 | 2.38 |
| VSEARCH | Only indels, short-long length | 98 | 8 | 1.00 | 1.29 |
| Velvet | Only indels, short-long length | 98 | 15 | 0.59 | 1.01 |
| Velvet | Only indels, short-long length | 94 | 15 | 0.59 | 1.01 |
| Velvet | Only indels, short-long length | 90 | 15 | 0.61 | 1.00 |

Table S2 – continued from previous page
| Assembler | Simulation | Percent match | K-mer length | Completeness | Contig/Fragment Ratio |
| --- | --- | --- | --- | --- | --- |
| Velvet | Only indels, short-long length | 98 | 31 | 0.77 | 0.83 |
| Velvet | Only indels, short-long length | 94 | 31 | 0.78 | 0.85 |
| Velvet | Only indels, short-long length | 90 | 31 | 0.80 | 0.89 |
| Stacks2 | Only indels, short-long length | 94 | 31 | 0.63 | 0.97 |
| Stacks2 | Only indels, short-long length | 98 | 31 | 0.64 | 0.99 |
| Stacks2 | Only indels, short-long length | 90 | opt | 0.62 | 0.96 |
| Stacks2 | Only indels, short-long length | 94 | 15 | 0.63 | 0.97 |
| Stacks2 | Only indels, short-long length | 98 | 15 | 0.64 | 0.99 |
| Stacks2 | Only indels, short-long length | 90 | 15 | 0.62 | 0.96 |
| Stacks2 | Only indels, short-long length | 94 | opt | 0.63 | 0.97 |
| Stacks2 | Only indels, short-long length | 98 | opt | 0.64 | 0.99 |
| Stacks2 | Only indels, short-long length | 90 | 31 | 0.62 | 0.97 |

**Table S3:** Assembler results for all simulations derived from the *H. sapiens* genome. Simulations were used to explore the impact of the number of mutations (overall and per locus), the proportion of mutations that were indels and the size of indels, on assembly performance across five different assemblers (i.e., CD-HIT, Stacks, Stacks2, VSEARCH, and Velvet), with varying percent match and k-mer lengths (when appropriate). Detailed descriptions of each simulation can be found in Table 1 of the main text. Assembly results were compared using two different metrics, the proportion of contigs that match genome fragments (completeness) and over-under assembly ratio (contigs:fragments). The completeness proportion includes exact sequence matches as well as close matches (see main text methods for clarification). K-mer length is left blank in the case of CD-HIT entries, because CD-HIT does not use k-mer length as a parameter setting.

| Assembler | Simulation | Percent match | K-mer length | Completeness | Contig/Fragment Ratio |
| --- | --- | --- | --- | --- | --- |
| CD-HIT | Original Genome | 94 | - | 0.92 | 0.92 |
| CD-HIT | Original Genome | 98 | - | 0.98 | 0.98 |
| CD-HIT | Original Genome | 90 | - | 0.88 | 0.88 |
| Stacks | Original Genome | 90 | 15 | 0.96 | 0.96 |
| Stacks | Original Genome | 98 | 15 | 0.96 | 0.97 |
| Stacks | Original Genome | 94 | 15 | 0.96 | 0.96 |
| Stacks | Original Genome | 90 | 31 | 0.96 | 0.96 |
| Stacks | Original Genome | 98 | 31 | 0.96 | 0.97 |
| Stacks | Original Genome | 94 | 31 | 0.96 | 0.96 |
| Stacks | Original Genome | 90 | opt | 0.96 | 0.96 |
| Stacks | Original Genome | 98 | opt | 0.96 | 0.97 |
| Stacks | Original Genome | 94 | opt | 0.96 | 0.96 |
| VSEARCH | Original Genome | 90 | 15 | 0.88 | 3.98 |
| VSEARCH | Original Genome | 90 | 8 | 0.88 | 3.52 |
| VSEARCH | Original Genome | 94 | 15 | 0.92 | 4.03 |
| VSEARCH | Original Genome | 94 | 8 | 0.92 | 3.56 |
| VSEARCH | Original Genome | 98 | 15 | 0.98 | 4.09 |
| VSEARCH | Original Genome | 98 | 8 | 0.98 | 3.62 |
| Velvet | Original Genome | 98 | 15 | 0.17 | 0.93 |
| Velvet | Original Genome | 94 | 15 | 0.17 | 0.93 |
| Velvet | Original Genome | 90 | 15 | 0.17 | 0.91 |
| Velvet | Original Genome | 98 | 31 | 0.76 | 0.77 |
| Velvet | Original Genome | 94 | 31 | 0.76 | 0.77 |
| Velvet | Original Genome | 90 | 31 | 0.76 | 0.77 |
| Stacks2 | Original Genome | 94 | 15 | 0.92 | 0.93 |
| Stacks2 | Original Genome | 94 | 31 | 0.91 | 0.92 |
| Stacks2 | Original Genome | 98 | 15 | 0.95 | 0.96 |
| Stacks2 | Original Genome | 98 | opt | 0.96 | 0.96 |
| Stacks2 | Original Genome | 94 | opt | 0.92 | 0.92 |
| Stacks2 | Original Genome | 90 | opt | 0.91 | 0.92 |
| Stacks2 | Original Genome | 90 | 31 | 0.89 | 0.91 |
| Stacks2 | Original Genome | 90 | 15 | 0.92 | 0.93 |
| Stacks2 | Original Genome | 98 | 31 | 0.95 | 0.96 |
| CD-HIT | A few SNPs | 94 | - | 0.92 | 0.92 |
| CD-HIT | A few SNPs | 98 | - | 0.98 | 1.02 |

Table S3 – continued from previous page
| Assembler | Simulation | Percent match | K-mer length | Completeness | Contig/Fragment Ratio |
| --- | --- | --- | --- | --- | --- |
| CD-HIT | A few SNPs | 90 | - | 0.87 | 0.87 |
| Stacks | A few SNPs | 90 | 15 | 0.94 | 0.95 |
| Stacks | A few SNPs | 98 | 15 | 0.95 | 0.96 |
| Stacks | A few SNPs | 94 | 15 | 0.95 | 0.95 |
| Stacks | A few SNPs | 90 | 31 | 0.95 | 0.95 |
| Stacks | A few SNPs | 98 | 31 | 0.95 | 0.96 |
| Stacks | A few SNPs | 94 | 31 | 0.95 | 0.95 |
| Stacks | A few SNPs | 90 | opt | 0.94 | 0.95 |
| Stacks | A few SNPs | 98 | opt | 0.95 | 0.96 |
| Stacks | A few SNPs | 94 | opt | 0.95 | 0.95 |
| VSEARCH | A few SNPs | 90 | 15 | 0.87 | 3.33 |
| VSEARCH | A few SNPs | 90 | 8 | 0.87 | 2.87 |
| VSEARCH | A few SNPs | 94 | 15 | 0.92 | 3.38 |
| VSEARCH | A few SNPs | 94 | 8 | 0.91 | 2.92 |
| VSEARCH | A few SNPs | 98 | 15 | 0.98 | 3.48 |
| VSEARCH | A few SNPs | 98 | 8 | 0.98 | 3.03 |
| Velvet | A few SNPs | 98 | 15 | 0.12 | 0.96 |
| Velvet | A few SNPs | 94 | 15 | 0.12 | 0.96 |
| Velvet | A few SNPs | 90 | 15 | 0.14 | 0.91 |
| Velvet | A few SNPs | 98 | 31 | 0.65 | 0.67 |
| Velvet | A few SNPs | 94 | 31 | 0.72 | 0.73 |
| Velvet | A few SNPs | 90 | 31 | 0.72 | 0.73 |
| Stacks2 | A few SNPs | 90 | 31 | 0.87 | 0.89 |
| Stacks2 | A few SNPs | 94 | 15 | 0.89 | 0.90 |
| Stacks2 | A few SNPs | 90 | opt | 0.86 | 0.88 |
| Stacks2 | A few SNPs | 98 | 15 | 0.94 | 0.95 |
| Stacks2 | A few SNPs | 98 | opt | 0.94 | 0.95 |
| Stacks2 | A few SNPs | 94 | 31 | 0.89 | 0.90 |
| Stacks2 | A few SNPs | 98 | 31 | 0.94 | 0.95 |
| Stacks2 | A few SNPs | 94 | opt | 0.89 | 0.90 |
| Stacks2 | A few SNPs | 90 | 15 | 0.86 | 0.88 |
| CD-HIT | More SNPs | 94 | - | 0.92 | 0.92 |
| CD-HIT | More SNPs | 98 | - | 0.98 | 1.02 |
| CD-HIT | More SNPs | 90 | - | 0.87 | 0.87 |
| Stacks | More SNPs | 90 | 15 | 0.94 | 0.95 |
| Stacks | More SNPs | 98 | 15 | 0.95 | 0.96 |
| Stacks | More SNPs | 94 | 15 | 0.95 | 0.95 |
| Stacks | More SNPs | 90 | 31 | 0.95 | 0.95 |
| Stacks | More SNPs | 98 | 31 | 0.95 | 0.96 |
| Stacks | More SNPs | 94 | 31 | 0.95 | 0.95 |
| Stacks | More SNPs | 90 | opt | 0.94 | 0.95 |
| Stacks | More SNPs | 98 | opt | 0.95 | 0.96 |
| Stacks | More SNPs | 94 | opt | 0.95 | 0.95 |
| VSEARCH | More SNPs | 90 | 15 | 0.87 | 3.88 |
| VSEARCH | More SNPs | 90 | 8 | 0.87 | 3.49 |
| VSEARCH | More SNPs | 94 | 15 | 0.92 | 3.93 |
| VSEARCH | More SNPs | 94 | 8 | 0.92 | 3.54 |
| VSEARCH | More SNPs | 98 | 15 | 0.98 | 4.03 |
| VSEARCH | More SNPs | 98 | 8 | 0.98 | 3.64 |
| Velvet | More SNPs | 98 | 15 | 0.11 | 0.96 |

Table S3 – continued from previous page
| Assembler | Simulation | Percent match | K-mer length | Completeness | Contig/Fragment Ratio |
| --- | --- | --- | --- | --- | --- |
| Velvet | More SNPs | 94 | 15 | 0.11 | 0.96 |
| Velvet | More SNPs | 90 | 15 | 0.14 | 0.91 |
| Velvet | More SNPs | 98 | 31 | 0.65 | 0.66 |
| Velvet | More SNPs | 94 | 31 | 0.72 | 0.73 |
| Velvet | More SNPs | 90 | 31 | 0.72 | 0.73 |
| Stacks2 | More SNPs | 90 | 31 | 0.87 | 0.89 |
| Stacks2 | More SNPs | 94 | 15 | 0.89 | 0.90 |
| Stacks2 | More SNPs | 98 | 31 | 0.94 | 0.95 |
| Stacks2 | More SNPs | 94 | 31 | 0.89 | 0.90 |
| Stacks2 | More SNPs | 98 | opt | 0.94 | 0.95 |
| Stacks2 | More SNPs | 90 | opt | 0.86 | 0.88 |
| Stacks2 | More SNPs | 98 | 15 | 0.94 | 0.95 |
| Stacks2 | More SNPs | 94 | opt | 0.89 | 0.89 |
| Stacks2 | More SNPs | 90 | 15 | 0.86 | 0.88 |
| CD-HIT | Multiple SNPs per locus | 94 | - | 0.92 | 0.92 |
| CD-HIT | Multiple SNPs per locus | 98 | - | 0.98 | 1.08 |
| CD-HIT | Multiple SNPs per locus | 90 | - | 0.87 | 0.87 |
| Stacks | Multiple SNPs per locus | 90 | 15 | 0.94 | 0.95 |
| Stacks | Multiple SNPs per locus | 98 | 15 | 0.95 | 0.97 |
| Stacks | Multiple SNPs per locus | 94 | 15 | 0.94 | 0.95 |
| Stacks | Multiple SNPs per locus | 90 | 31 | 0.94 | 0.95 |
| Stacks | Multiple SNPs per locus | 98 | 31 | 0.95 | 0.97 |
| Stacks | Multiple SNPs per locus | 94 | 31 | 0.94 | 0.95 |
| Stacks | Multiple SNPs per locus | 90 | opt | 0.94 | 0.95 |
| Stacks | Multiple SNPs per locus | 98 | opt | 0.95 | 0.97 |
| Stacks | Multiple SNPs per locus | 94 | opt | 0.94 | 0.95 |
| VSEARCH | Multiple SNPs per locus | 90 | 15 | 0.87 | 3.91 |
| VSEARCH | Multiple SNPs per locus | 90 | 8 | 0.87 | 3.45 |
| VSEARCH | Multiple SNPs per locus | 94 | 15 | 0.92 | 3.96 |
| VSEARCH | Multiple SNPs per locus | 94 | 8 | 0.92 | 3.50 |
| VSEARCH | Multiple SNPs per locus | 98 | 15 | 0.98 | 4.12 |
| VSEARCH | Multiple SNPs per locus | 98 | 8 | 0.98 | 3.66 |
| Velvet | Multiple SNPs per locus | 98 | 15 | 0.10 | 0.97 |
| Velvet | Multiple SNPs per locus | 94 | 15 | 0.10 | 0.97 |
| Velvet | Multiple SNPs per locus | 90 | 15 | 0.13 | 0.91 |
| Velvet | Multiple SNPs per locus | 98 | 31 | 0.62 | 0.64 |
| Velvet | Multiple SNPs per locus | 94 | 31 | 0.71 | 0.72 |
| Velvet | Multiple SNPs per locus | 90 | 31 | 0.71 | 0.73 |
| Stacks2 | Multiple SNPs per locus | 98 | 31 | 0.94 | 0.96 |
| Stacks2 | Multiple SNPs per locus | 90 | 15 | 0.86 | 0.87 |
| Stacks2 | Multiple SNPs per locus | 98 | opt | 0.94 | 0.96 |
| Stacks2 | Multiple SNPs per locus | 90 | opt | 0.86 | 0.87 |
| Stacks2 | Multiple SNPs per locus | 94 | 31 | 0.89 | 0.90 |
| Stacks2 | Multiple SNPs per locus | 90 | 31 | 0.87 | 0.89 |
| Stacks2 | Multiple SNPs per locus | 94 | 15 | 0.88 | 0.89 |
| Stacks2 | Multiple SNPs per locus | 94 | opt | 0.89 | 0.90 |
| Stacks2 | Multiple SNPs per locus | 98 | 15 | 0.94 | 0.96 |
| CD-HIT | Mostly SNPs + few indels | 94 | - | 0.92 | 0.92 |
| CD-HIT | Mostly SNPs + few indels | 98 | - | 0.98 | 1.03 |
| CD-HIT | Mostly SNPs + few indels | 90 | - | 0.87 | 0.87 |

Table S3 – continued from previous page
| Assembler | Simulation | Percent match | K-mer length | Completeness | Contig/Fragment Ratio |
| --- | --- | --- | --- | --- | --- |
| Stacks | Mostly SNPs + few indels | 90 | 15 | 0.95 | 0.98 |
| Stacks | Mostly SNPs + few indels | 98 | 15 | 0.96 | 1.00 |
| Stacks | Mostly SNPs + few indels | 94 | 15 | 0.95 | 0.99 |
| Stacks | Mostly SNPs + few indels | 90 | 31 | 0.95 | 0.98 |
| Stacks | Mostly SNPs + few indels | 98 | 31 | 0.95 | 1.00 |
| Stacks | Mostly SNPs + few indels | 94 | 31 | 0.95 | 0.99 |
| Stacks | Mostly SNPs + few indels | 90 | opt | 0.95 | 0.98 |
| Stacks | Mostly SNPs + few indels | 98 | opt | 0.96 | 1.00 |
| Stacks | Mostly SNPs + few indels | 94 | opt | 0.95 | 0.99 |
| VSEARCH | Mostly SNPs + few indels | 90 | 15 | 0.87 | 3.89 |
| VSEARCH | Mostly SNPs + few indels | 90 | 8 | 0.87 | 3.51 |
| VSEARCH | Mostly SNPs + few indels | 94 | 15 | 0.92 | 3.93 |
| VSEARCH | Mostly SNPs + few indels | 94 | 8 | 0.92 | 3.56 |
| VSEARCH | Mostly SNPs + few indels | 98 | 15 | 0.98 | 4.04 |
| VSEARCH | Mostly SNPs + few indels | 98 | 8 | 0.98 | 3.67 |
| Velvet | Mostly SNPs + few indels | 98 | 15 | 0.11 | 0.96 |
| Velvet | Mostly SNPs + few indels | 94 | 15 | 0.11 | 0.96 |
| Velvet | Mostly SNPs + few indels | 90 | 15 | 0.14 | 0.91 |
| Velvet | Mostly SNPs + few indels | 98 | 31 | 0.65 | 0.67 |
| Velvet | Mostly SNPs + few indels | 94 | 31 | 0.72 | 0.73 |
| Velvet | Mostly SNPs + few indels | 90 | 31 | 0.72 | 0.73 |
| Stacks2 | Mostly SNPs + few indels | 90 | opt | 0.83 | 0.88 |
| Stacks2 | Mostly SNPs + few indels | 94 | opt | 0.85 | 0.90 |
| Stacks2 | Mostly SNPs + few indels | 98 | opt | 0.90 | 0.95 |
| Stacks2 | Mostly SNPs + few indels | 94 | 15 | 0.85 | 0.90 |
| Stacks2 | Mostly SNPs + few indels | 98 | 31 | 0.90 | 0.95 |
| Stacks2 | Mostly SNPs + few indels | 98 | 15 | 0.90 | 0.95 |
| Stacks2 | Mostly SNPs + few indels | 94 | 31 | 0.86 | 0.90 |
| Stacks2 | Mostly SNPs + few indels | 90 | 31 | 0.84 | 0.89 |
| Stacks2 | Mostly SNPs + few indels | 90 | 15 | 0.83 | 0.88 |
| CD-HIT | 50:50 SNPs:indels | 94 | - | 0.92 | 0.92 |
| CD-HIT | 50:50 SNPs:indels | 98 | - | 0.98 | 1.02 |
| CD-HIT | 50:50 SNPs:indels | 90 | - | 0.87 | 0.87 |
| Stacks | 50:50 SNPs:indels | 90 | 15 | 0.95 | 1.09 |
| Stacks | 50:50 SNPs:indels | 98 | 15 | 0.96 | 1.14 |
| Stacks | 50:50 SNPs:indels | 94 | 15 | 0.95 | 1.11 |
| Stacks | 50:50 SNPs:indels | 90 | 31 | 0.95 | 1.10 |
| Stacks | 50:50 SNPs:indels | 98 | 31 | 0.96 | 1.14 |
| Stacks | 50:50 SNPs:indels | 94 | 31 | 0.95 | 1.11 |
| Stacks | 50:50 SNPs:indels | 90 | opt | 0.95 | 1.09 |
| Stacks | 50:50 SNPs:indels | 98 | opt | 0.96 | 1.14 |
| Stacks | 50:50 SNPs:indels | 94 | opt | 0.95 | 1.11 |
| VSEARCH | 50:50 SNPs:indels | 90 | 15 | 0.87 | 3.91 |
| VSEARCH | 50:50 SNPs:indels | 90 | 8 | 0.87 | 3.45 |
| VSEARCH | 50:50 SNPs:indels | 94 | 15 | 0.92 | 3.96 |
| VSEARCH | 50:50 SNPs:indels | 94 | 8 | 0.92 | 3.50 |
| VSEARCH | 50:50 SNPs:indels | 98 | 15 | 0.98 | 4.07 |
| VSEARCH | 50:50 SNPs:indels | 98 | 8 | 0.98 | 3.62 |
| Velvet | 50:50 SNPs:indels | 98 | 15 | 0.11 | 0.93 |
| Velvet | 50:50 SNPs:indels | 94 | 15 | 0.11 | 0.93 |

Table S3 – continued from previous page
| Assembler | Simulation | Percent match | K-mer length | Completeness | Contig/Fragment Ratio |
| --- | --- | --- | --- | --- | --- |
| Velvet | 50:50 SNPs:indels | 90 | 15 | 0.13 | 0.91 |
| Velvet | 50:50 SNPs:indels | 98 | 31 | 0.63 | 0.67 |
| Velvet | 50:50 SNPs:indels | 94 | 31 | 0.68 | 0.73 |
| Velvet | 50:50 SNPs:indels | 90 | 31 | 0.69 | 0.73 |
| Stacks2 | 50:50 SNPs:indels | 98 | opt | 0.77 | 0.95 |
| Stacks2 | 50:50 SNPs:indels | 98 | 15 | 0.77 | 0.95 |
| Stacks2 | 50:50 SNPs:indels | 94 | 15 | 0.73 | 0.90 |
| Stacks2 | 50:50 SNPs:indels | 90 | 15 | 0.71 | 0.89 |
| Stacks2 | 50:50 SNPs:indels | 90 | opt | 0.71 | 0.89 |
| Stacks2 | 50:50 SNPs:indels | 98 | 31 | 0.77 | 0.95 |
| Stacks2 | 50:50 SNPs:indels | 94 | 31 | 0.73 | 0.91 |
| Stacks2 | 50:50 SNPs:indels | 94 | opt | 0.73 | 0.91 |
| Stacks2 | 50:50 SNPs:indels | 90 | 31 | 0.72 | 0.90 |
| CD-HIT | Only indels | 94 | - | 0.92 | 0.92 |
| CD-HIT | Only indels | 98 | - | 0.98 | 1.01 |
| CD-HIT | Only indels | 90 | - | 0.87 | 0.87 |
| Stacks | Only indels | 90 | 15 | 0.96 | 1.23 |
| Stacks | Only indels | 98 | 15 | 0.97 | 1.30 |
| Stacks | Only indels | 94 | 15 | 0.96 | 1.26 |
| Stacks | Only indels | 90 | 31 | 0.96 | 1.23 |
| Stacks | Only indels | 98 | 31 | 0.97 | 1.30 |
| Stacks | Only indels | 94 | 31 | 0.96 | 1.26 |
| Stacks | Only indels | 90 | opt | 0.96 | 1.23 |
| Stacks | Only indels | 98 | opt | 0.97 | 1.30 |
| Stacks | Only indels | 94 | opt | 0.96 | 1.26 |
| VSEARCH | Only indels | 90 | 15 | 0.87 | 3.83 |
| VSEARCH | Only indels | 90 | 8 | 0.87 | 3.41 |
| VSEARCH | Only indels | 94 | 15 | 0.92 | 3.88 |
| VSEARCH | Only indels | 94 | 8 | 0.92 | 3.46 |
| VSEARCH | Only indels | 98 | 15 | 0.98 | 3.99 |
| VSEARCH | Only indels | 98 | 8 | 0.98 | 3.58 |
| Velvet | Only indels | 98 | 15 | 0.11 | 0.91 |
| Velvet | Only indels | 94 | 15 | 0.11 | 0.91 |
| Velvet | Only indels | 90 | 15 | 0.12 | 0.90 |
| Velvet | Only indels | 98 | 31 | 0.61 | 0.66 |
| Velvet | Only indels | 94 | 31 | 0.65 | 0.73 |
| Velvet | Only indels | 90 | 31 | 0.65 | 0.73 |
| Stacks2 | Only indels | 90 | 31 | 0.58 | 0.91 |
| Stacks2 | Only indels | 94 | 15 | 0.59 | 0.92 |
| Stacks2 | Only indels | 94 | 31 | 0.59 | 0.92 |
| Stacks2 | Only indels | 94 | opt | 0.59 | 0.92 |
| Stacks2 | Only indels | 98 | 31 | 0.62 | 0.96 |
| Stacks2 | Only indels | 98 | opt | 0.62 | 0.96 |
| Stacks2 | Only indels | 90 | 15 | 0.57 | 0.90 |
| Stacks2 | Only indels | 98 | 15 | 0.62 | 0.96 |
| Stacks2 | Only indels | 90 | opt | 0.57 | 0.90 |
| CD-HIT | Only indels, short-mid length | 94 | - | 0.92 | 0.93 |
| CD-HIT | Only indels, short-mid length | 98 | - | 0.99 | 1.22 |
| CD-HIT | Only indels, short-mid length | 90 | - | 0.87 | 0.87 |
| Stacks | Only indels, short-mid length | 90 | 15 | 0.96 | 1.23 |

Table S3 – continued from previous page
| Assembler | Simulation | Percent match | K-mer length | Completeness | Contig/Fragment Ratio |
| --- | --- | --- | --- | --- | --- |
| Stacks | Only indels, short-mid length | 98 | 15 | 0.97 | 1.30 |
| Stacks | Only indels, short-mid length | 94 | 15 | 0.96 | 1.26 |
| Stacks | Only indels, short-mid length | 90 | 31 | 0.96 | 1.24 |
| Stacks | Only indels, short-mid length | 98 | 31 | 0.97 | 1.30 |
| Stacks | Only indels, short-mid length | 94 | 31 | 0.96 | 1.26 |
| Stacks | Only indels, short-mid length | 90 | opt | 0.96 | 1.23 |
| Stacks | Only indels, short-mid length | 98 | opt | 0.97 | 1.30 |
| Stacks | Only indels, short-mid length | 94 | opt | 0.96 | 1.26 |
| VSEARCH | Only indels, short-mid length | 90 | 15 | 0.88 | 3.92 |
| VSEARCH | Only indels, short-mid length | 90 | 8 | 0.87 | 3.50 |
| VSEARCH | Only indels, short-mid length | 94 | 15 | 0.92 | 3.98 |
| VSEARCH | Only indels, short-mid length | 94 | 8 | 0.92 | 3.56 |
| VSEARCH | Only indels, short-mid length | 98 | 15 | 0.98 | 4.27 |
| VSEARCH | Only indels, short-mid length | 98 | 8 | 0.98 | 3.85 |
| Velvet | Only indels, short-mid length | 98 | 15 | 0.11 | 0.93 |
| Velvet | Only indels, short-mid length | 94 | 15 | 0.11 | 0.93 |
| Velvet | Only indels, short-mid length | 90 | 15 | 0.12 | 0.92 |
| Velvet | Only indels, short-mid length | 98 | 31 | 0.61 | 0.67 |
| Velvet | Only indels, short-mid length | 94 | 31 | 0.63 | 0.69 |
| Velvet | Only indels, short-mid length | 90 | 31 | 0.65 | 0.73 |
| Stacks2 | Only indels, short-mid length | 90 | opt | 0.58 | 0.90 |
| Stacks2 | Only indels, short-mid length | 98 | 31 | 0.62 | 0.96 |
| Stacks2 | Only indels, short-mid length | 94 | 31 | 0.59 | 0.92 |
| Stacks2 | Only indels, short-mid length | 90 | 15 | 0.57 | 0.90 |
| Stacks2 | Only indels, short-mid length | 90 | 31 | 0.58 | 0.91 |
| Stacks2 | Only indels, short-mid length | 98 | opt | 0.63 | 0.96 |
| Stacks2 | Only indels, short-mid length | 94 | opt | 0.59 | 0.92 |
| Stacks2 | Only indels, short-mid length | 94 | 15 | 0.59 | 0.91 |
| Stacks2 | Only indels, short-mid length | 98 | 15 | 0.63 | 0.96 |
| CD-HIT | Only indels, short-long length | 94 | - | 0.93 | 0.95 |
| CD-HIT | Only indels, short-long length | 98 | - | 0.99 | 1.26 |
| CD-HIT | Only indels, short-long length | 90 | - | 0.88 | 0.88 |
| Stacks | Only indels, short-long length | 90 | 15 | 0.96 | 1.23 |
| Stacks | Only indels, short-long length | 98 | 15 | 0.98 | 1.30 |
| Stacks | Only indels, short-long length | 94 | 15 | 0.96 | 1.27 |
| Stacks | Only indels, short-long length | 90 | 31 | 0.96 | 1.24 |
| Stacks | Only indels, short-long length | 98 | 31 | 0.98 | 1.30 |
| Stacks | Only indels, short-long length | 94 | 31 | 0.96 | 1.27 |
| Stacks | Only indels, short-long length | 90 | opt | 0.96 | 1.23 |
| Stacks | Only indels, short-long length | 98 | opt | 0.98 | 1.30 |
| Stacks | Only indels, short-long length | 94 | opt | 0.96 | 1.27 |
| VSEARCH | Only indels, short-long length | 90 | 15 | 0.88 | 3.99 |
| VSEARCH | Only indels, short-long length | 90 | 8 | 0.88 | 3.52 |
| VSEARCH | Only indels, short-long length | 94 | 15 | 0.93 | 4.06 |
| VSEARCH | Only indels, short-long length | 94 | 8 | 0.93 | 3.59 |
| VSEARCH | Only indels, short-long length | 98 | 15 | 0.99 | 4.37 |
| VSEARCH | Only indels, short-long length | 98 | 8 | 0.99 | 3.91 |
| Velvet | Only indels, short-long length | 98 | 15 | 0.11 | 0.94 |
| Velvet | Only indels, short-long length | 94 | 15 | 0.11 | 0.94 |
| Velvet | Only indels, short-long length | 90 | 15 | 0.12 | 0.93 |

Table S3 – continued from previous page
| Assembler | Simulation | Percent match | K-mer length | Completeness | Contig/Fragment Ratio |
| --- | --- | --- | --- | --- | --- |
| Velvet | Only indels, short-long length | 98 | 31 | 0.62 | 0.67 |
| Velvet | Only indels, short-long length | 94 | 31 | 0.62 | 0.69 |
| Velvet | Only indels, short-long length | 90 | 31 | 0.64 | 0.71 |
| Stacks2 | Only indels, short-long length | 98 | opt | 0.63 | 0.97 |
| Stacks2 | Only indels, short-long length | 94 | 31 | 0.59 | 0.92 |
| Stacks2 | Only indels, short-long length | 90 | opt | 0.58 | 0.90 |
| Stacks2 | Only indels, short-long length | 98 | 15 | 0.63 | 0.97 |
| Stacks2 | Only indels, short-long length | 90 | 15 | 0.58 | 0.90 |
| Stacks2 | Only indels, short-long length | 90 | 31 | 0.58 | 0.91 |
| Stacks2 | Only indels, short-long length | 94 | 15 | 0.59 | 0.92 |
| Stacks2 | Only indels, short-long length | 94 | opt | 0.59 | 0.92 |
| Stacks2 | Only indels, short-long length | 98 | 31 | 0.63 | 0.97 |

